# The adaptive benefit of evolved increases in hemoglobin-O_2_ affinity is contingent on tissue O_2_ diffusing capacity in high-altitude deer mice

**DOI:** 10.1101/2020.10.29.357665

**Authors:** Oliver H. Wearing, Catherine M. Ivy, Natalia Gutiérrez-Pinto, Jonathan P. Velotta, Shane C. Campbell-Staton, Chandrasekhar Natarajan, Zachary A. Cheviron, Jay F. Storz, Graham R. Scott

**Affiliations:** Department of Biology, McMaster University, Hamilton, ON L8S 4K1, Canada; School of Biological Sciences, University of Nebraska, Lincoln, NE 68588, USA; Department of Biological Sciences, University of Denver, Denver CO 80210, USA; Department of Ecology and Evolutionary Biology, Institute for Society and Genetics, University of California, Los Angeles, CA 90095, USA; Division of Biological Sciences, University of Montana, Missoula, MT 59812

**Keywords:** Evolutionary physiology, high-altitude adaptation, O_2_ transport pathway, complex trait evolution, hemoglobin adaptation

## Abstract

**Background:** Complex organismal traits are often the result of multiple interacting genes and sub-organismal phenotypes, but how these interactions shape the evolutionary trajectories of adaptive traits is poorly understood. We examined how functional interactions between cardiorespiratory traits contribute to adaptive increases in the capacity for aerobic thermogenesis (maximal O_2_ consumption, V◻O_2_max, during acute cold exposure) in high-altitude deer mice (*Peromyscus maniculatus*). We crossed highland and lowland deer mice to produce F_2_ inter-population hybrids, which expressed genetically based variation in hemoglobin (Hb) O_2_ affinity on a mixed genetic background. We then combined physiological experiments and mathematical modeling of the O_2_ transport pathway to examine links between cardiorespiratory traits and V◻O_2_max.

**Results:** Physiological experiments revealed that increases in Hb-O_2_ affinity of red blood cells improved blood oxygenation in hypoxia, but were not associated with an enhancement in V◻O_2_max. Sensitivity analyses performed using mathematical modeling showed that the influence of Hb-O_2_ affinity on V◻O_2_max in hypoxia was contingent on the capacity for O_2_ diffusion in active tissues.

**Conclusions:** These results suggest that increases in Hb-O_2_ affinity would only have adaptive value in hypoxic conditions if concurrent with or preceded by increases in tissue O_2_ diffusing capacity. In high-altitude deer mice, the adaptive benefit of increasing Hb-O_2_ affinity is contingent on the capacity to extract O_2_ from the blood, which helps resolve controversies about the general role of hemoglobin function in hypoxia tolerance.

## Background

A long-standing goal of evolutionary biology is to understand how the functional integration of traits influences patterns of phenotypic change and adaptation (1). Complex physiological phenotypes often represent an emergent property of functional interactions among different tissues and organ systems, which in turn may be developmentally interrelated and genetically correlated. The functional, developmental, and genetic interdependence of traits may facilitate environmental adaptation if semi-autonomous components of a complex phenotype respond synergistically to selection. Alternatively, functional integration and genetic correlations among components of a trait can limit and channel pathways of phenotypic evolution (2, 3). Evolutionary questions about phenotypic integration and adaptation can be addressed most profitably by examining well-defined traits with well-characterized functions and well-documented associations with fitness under natural conditions.

The capacity for aerobic thermogenesis in small mammals at high altitude is a complex performance trait that is well suited to experimental studies of how patterns of phenotypic integration affect the process of adaptation. At high altitude, cold temperatures challenge the ability of endotherms to maintain body temperature and activity, which is especially difficult in smaller animals that have high surface area to volume ratios. Unsurprisingly, aerobic thermogenesis (quantified as maximal O_2_ consumption, V◻O_2_max, during acute cold exposure) in hypoxia is under strong directional selection in some small mammals at high altitude (4), which have evolved higher thermogenic V◻O_2_max (5–9). Thermogenic V◻O_2_max is supported by the integrated function of the O_2_ transport pathway, the conceptual steps (ventilation, pulmonary diffusion, circulation, tissue diffusion, and mitochondrial O_2_ utilization) involved in transporting O_2_ from inspired air to thermogenic tissues where O_2_ is used by mitochondria to support oxidative phosphorylation (10, 11). Therefore, studies of thermogenic V◻O_2_max in high-altitude natives are ideal for understanding the mechanisms underlying the adaptive evolution of complex traits.

Evolved increases in the O_2_ affinity of hemoglobin (Hb) are pervasive in high-altitude taxa, and have become classic examples of biochemical adaptation (12). However, the nature of the direct adaptive benefit conferred by increases in Hb-O_2_ affinity in highland species is controversial. Many highland taxa have evolved increases in Hb-O_2_ affinity independently, and in many cases, the molecular mechanisms underlying these changes in protein function are documented in detail (12–18). These increases in Hb-O_2_ affinity are often presumed to safeguard arterial O_2_ saturation in hypoxia and thus help improve tissue O_2_ delivery and aerobic capacity (5, 6, 19–26), although this has rarely been tested. Nonetheless, the relationship between Hb-O_2_ affinity and V◻O_2_max in hypoxia remains contentious (12, 25, 27–29). Theoretical modeling of the O_2_ transport pathway in humans suggests that increases in Hb-O_2_ affinity do not increase aerobic capacity in hypoxia on their own (30), because the advantage of increasing Hb-O_2_ affinity may be offset by a trade-off in O_2_ offloading at tissues (11, 31, 32). A recent study in humans with rare genetic Hb variants found that increases in Hb-O_2_ affinity attenuated the hypoxia-induced decline in aerobic capacity, but subjects with high Hb-O_2_ affinity also had compensatory polycythemia (33). Considering the strong functional integration of Hb within the O_2_ transport pathway, the advantages of increasing Hb-O_2_ affinity in high-altitude taxa may be contingent on the evolution of other cardiorespiratory traits, but this has not been experimentally investigated.

We sought to determine the effects of evolved increases in Hb-O_2_ affinity in high-altitude deer mice (*Peromyscus maniculatus*) on thermogenic V◻O_2_max in hypoxia, and to examine whether the adaptive benefit of changes in Hb-O_2_ affinity is contingent on other cardiorespiratory changes. Deer mice have the broadest altitudinal range of any North American mammal (34), ranging from near sea level to montane environments up to approx. 4,350 m above sea level (35), and high-altitude populations have evolved elevated thermogenic V◻O_2_max in hypoxia in response to directional selection (4–9). In conjunction with a higher V◻O_2_max in chronic hypoxia, high-altitude deer mice also exhibit higher pulmonary O_2_ extraction, arterial O_2_ saturation, cardiac output, and tissue O_2_ extraction than their lowland counterparts (9, 36). The latter is associated with several evolved changes in skeletal muscle phenotype and mitochondrial function (37–41). Highlanders have also evolved a higher Hb-O_2_ affinity as a result of amino acid replacements in duplicated genes that encode the α- and β-chain subunits of the α_2_β_2_ Hb tetramer (5, 6, 15, 20, 21, 23, 24, 26, 34, 42). This evolved increase in Hb-O_2_ affinity in highlanders is not complemented by an enhanced Bohr effect to augment O_2_ unloading (43). We initially hypothesised that these evolved increases in Hb-O_2_ affinity would be responsible for higher thermogenic capacity in highland deer mice, compared to their lowland conspecifics. To investigate the effect of genetically based changes in Hb-O_2_ affinity on whole-animal performance in hypoxia, we created F_2_ hybrids between high- and low-altitude deer mice (F_2_ intercross breeding design) to randomize associations between allelic globin variants, and we then examined the effects of α- and β-globin variants on red blood cell *P*_50_ (the O_2_ pressure, *P*O_2_, at which Hb is 50% saturated), arterial O_2_ saturation, thermogenic V◻O_2_max, and other physiological traits on an admixed genetic background (see Additional file 1: Fig. S1 for graphical overview of experimental design). We performed physiological measurements before and after chronic exposure to hypoxia to test for effects of Hb genotype on trait-specific acclimation responses. We then used our empirical data in an *in silico* model of the O_2_ transport pathway to examine the interactive effects of Hb-O_2_ affinity and the O_2_ diffusing capacity of tissues (D_T_O_2_) on V◻O_2_max. Our results suggest that increases in Hb-O_2_ affinity only contribute to the adaptive enhancement of thermogenic V◻O_2_max in hypoxia if accompanied by a corresponding increase in D_T_O_2_ to augment tissue O_2_ extraction.

## Results

We measured thermogenic V◻O_2_max, arterial O_2_ saturation, and other cardiorespiratory traits *in vivo* during acute exposure to cold heliox in normoxia (21% O_2_) and hypoxia (12% O_2_) in male and female F_2_ hybrid mice (mean ± SEM of body mass before hypoxia acclimation, 23.8 ± 0.9 g; see Additional file 2: Fig. S2 for body masses for each genotype before and after hypoxia acclimation) that possessed a diverse array of different α- and β-globin genotypes. We also performed *in vitro* measurements of red blood cell *P*_50_ using erythrocyte suspensions from the same set of mice. The F_2_ hybrids were generated by crossing wild mice from populations at high and low altitudes to produce F_1_ inter-population hybrids, followed by full-sibling matings to create 4 families of F_2_ hybrid progeny with admixed genetic backgrounds. Measurements of physiological phenotypes were made before and after a 6-week acclimation period to hypobaric hypoxia (12 kPa O_2_, simulating ~4,300 m above sea level). In general, hypoxia acclimation was associated with increased V◻O_2_max in hypoxia, along with increases in pulmonary ventilation, arterial O_2_ saturation, heart rate, hematocrit (Hct) and blood Hb concentration ([Hb]), but also increases in red blood cell *P*_50_ (Fig. 1, Additional file 3: Tables S1 and S2). However, hypoxia acclimation did not affect V◻O_2_max under normoxic conditions. Below, we describe the effects of Hb genotype on thermogenic V◻O_2_max and hematological traits in mice acclimated to normoxia, and then we describe how Hb genotype affects acclimation responses to chronic hypoxia.

**Figure 1.**
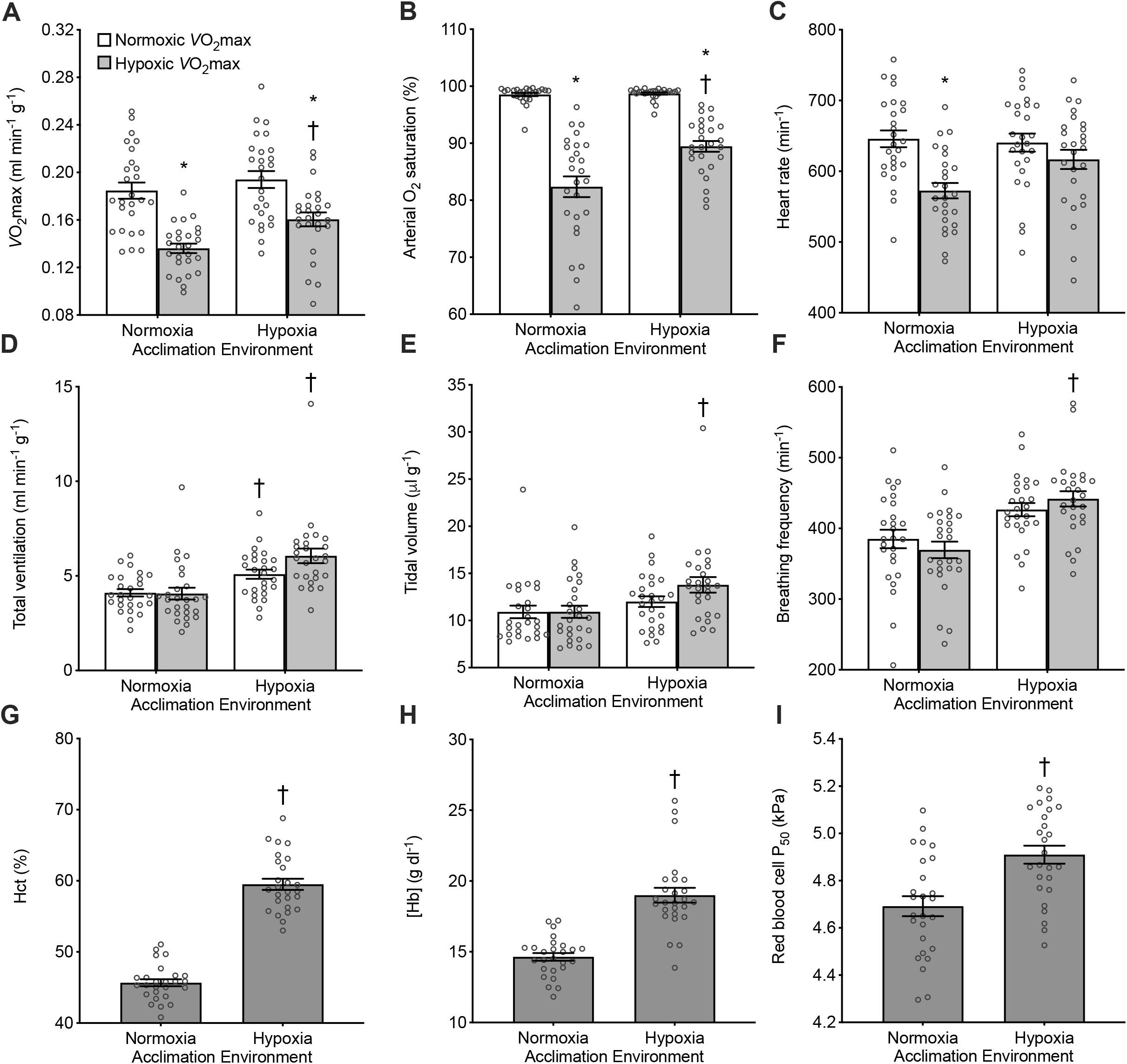
Acclimation responses to chronic hypoxia in all F_2_ inter-population hybrid deer mice of all genotypes for measurements at V◻O_2_max in normoxia (21 kPa O_2_) and hypoxia (12 kPa O_2_), and for measurements of hematology. Hct, hematocrit; [Hb], blood hemoglobin content; *P*_50_, O_2_ pressure at 50% saturation. *P < 0.05 between measurements in normoxia versus hypoxia within an acclimation condition. ^†^P < 0.05 vs. pre-acclimation value. Bars display mean ± SEM (n = 26) with individual data superimposed (circles).

### Genetically based decreases in red blood cell *P*_50_ improved arterial O_2_ saturation in hypoxia

In normoxia-acclimated mice, there was a significant main effect of Hb genotype on red blood cell *P*_50_ (P = 0.0048; Fig. 2A, Additional file 3: Table S3), which appeared to be largely attributable to the effects of α-globin variants. Mice possessing highland α-globin variants had a lower red blood cell *P*_50_ compared to those possessing lowland variants, reflecting a higher affinity for O_2_. In contrast, Hb genotype did not affect Hct (P = 0.8339), [Hb] (P = 0.9351), or the Hill coefficient (*n*) that quantifies the cooperativity of Hb-O_2_ binding (P = 0.8053; Additional file 4: Fig. S3, Additional file 3: Table S3).

**Figure 2.**
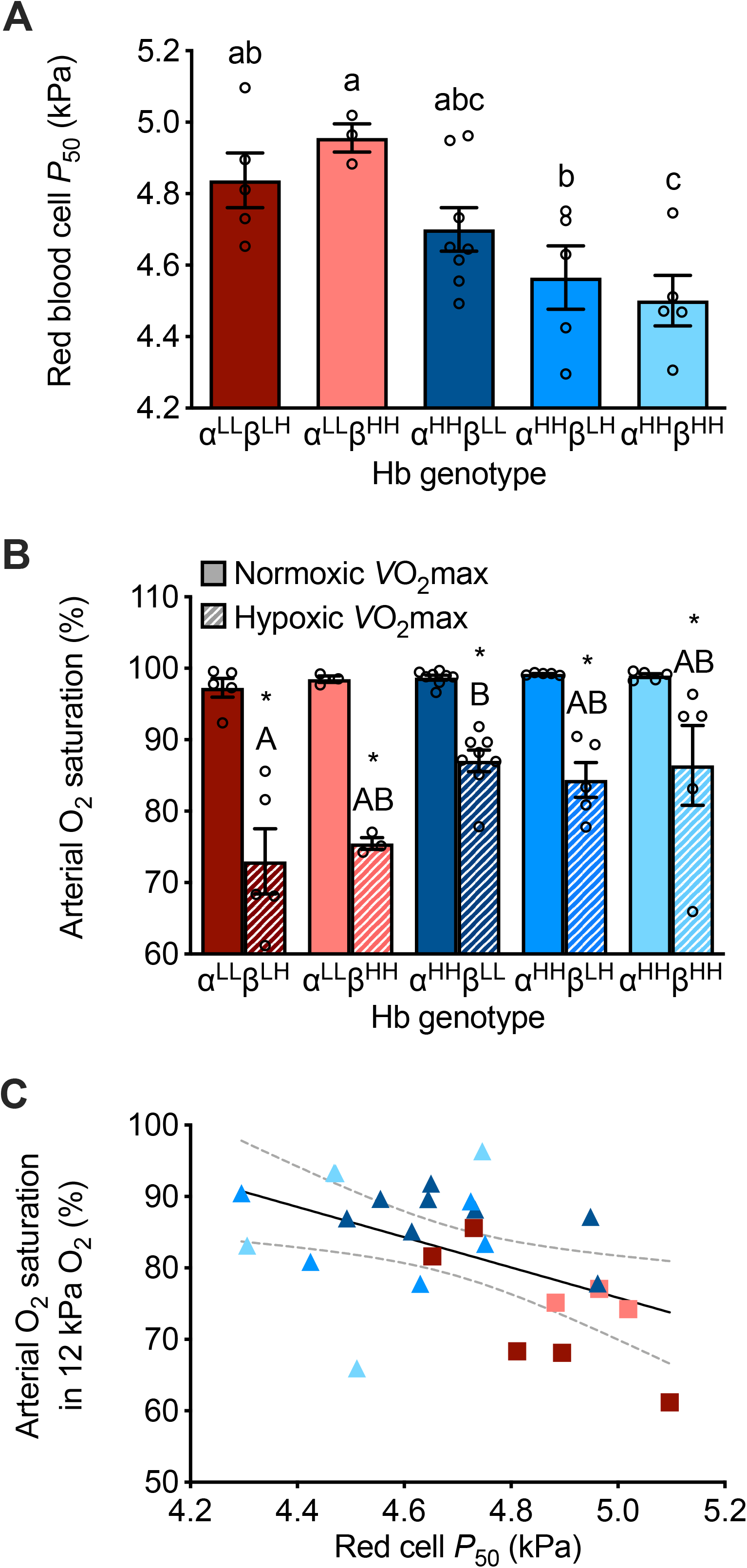
Variation in red blood cell O_2_ affinity and arterial O_2_ saturation associated with hemoglobin (Hb) genotype in F_2_ inter-population hybrid deer mice acclimated to normoxia. **A**) Red blood cell *P*_50_ (O_2_ pressure at 50% saturation). **B**) Arterial O_2_ saturation at V◻O_2_max measured in normoxia (21 kPa O_2_) and hypoxia (12 kPa O_2_). Bars display mean ± SEM (n = 3-8) with individual data superimposed (circles). Different α- and β-globin genotypes are shown as superscripts with ‘^L^’ representing the lowland haplotype and ‘^H^’ representing the highland haplotype. *P < 0.05, hypoxia vs. normoxia within a genotype. P < 0.05 between genotypes for values not sharing a letter. **C**) Linear regression of arterial O_2_ saturation in hypoxia and red blood cell *P*_50_ for individual data (P = 0.0103, R^2^ = 0.2441; dotted line represents 95% confidence interval). Symbol colors reflect Hb genotype, as shown in A and B.

Arterial O_2_ saturation varied in association with red blood cell *P*_50_ in hypoxia, but not in normoxia. There were significant main effects of both Hb genotype (P = 0.0189) and inspired *P*O_2_ (P < 0.0001) on arterial O_2_ saturation at V◻O_2_max, with mice exhibiting reduced saturation in hypoxia (Fig. 2B, Additional file 3: Table S3). However, the effect of inspired *P*O_2_ on arterial O_2_ saturation was influenced by genotype (genotype x *P*O_2_ interaction, P = 0.0389), as mice with the highland α-globin genotype exhibited a smaller reduction in arterial O_2_ saturation under hypoxia compared to those with the lowland genotype. Consequently, mice with highland α-globin maintained 9-14% higher arterial O_2_ saturation on average than those with lowland α-globin at hypoxic V◻O_2_max. Higher red blood cell O_2_ affinity was associated with higher arterial O_2_ saturation in hypoxia, as indicated by a significant negative relationship between arterial O_2_ saturation and red blood cell *P*_50_ (P = 0.0103, R^2^ = 0.2441; Fig. 2C).

### Genetically based variation in red blood cell *P*_50_ and arterial O_2_ saturation had no effect on thermogenicV◻O_2_max in hypoxia

V◻O_2_max was significantly reduced in hypoxia compared to normoxia by ~24% on average (P < 0.0001; Fig. 3A, Additional file 3: Table S3). However, although Hb genotype had a significant main effect on V◻O_2_max (P = 0.0416), V◻O_2_max in hypoxia did not follow the pattern of variation seen for arterial O_2_ saturation. As such, hypoxic V◻O_2_max was not correlated with arterial O_2_ saturation in hypoxia (Fig. 3B). Instead, the observed variation in V◻O_2_max appeared to be associated with variation in heart rate, which was also significantly affected by inspired *P*O_2_ (P < 0.0001), though the effect of genotype was only marginally significant (P = 0.0545; Additional file 5: Fig. S4A, Additional file 3: Table S3). Total ventilation, tidal volume, and breathing frequency were unaffected by Hb genotype (Additional file 5: Fig. S4, Additional file 3: Table S3).

**Figure 3.**
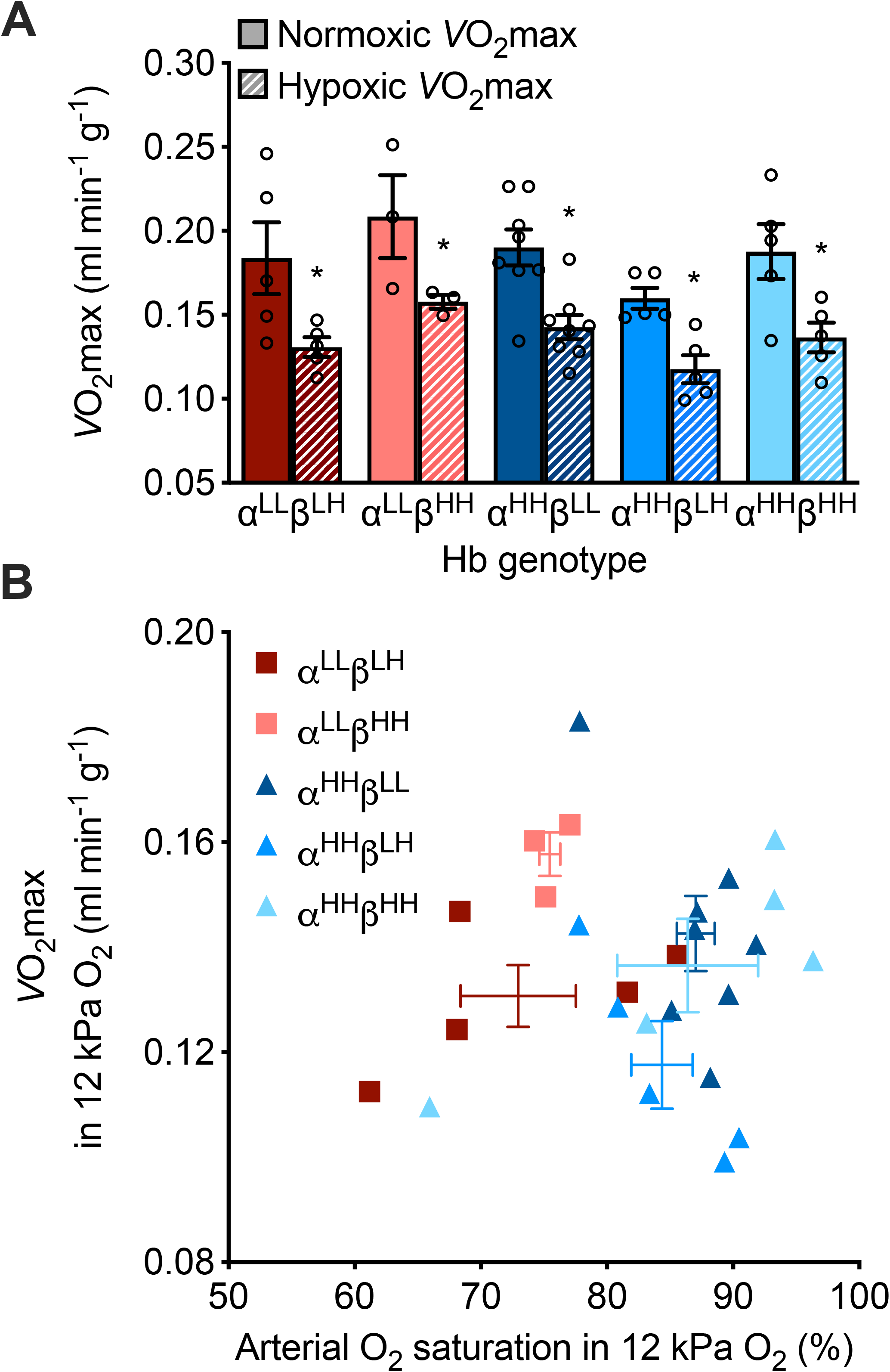
Variation in V◻O_2_max was unrelated to variation in arterial O_2_ saturation in F_2_ inter-population hybrid deer mice acclimated to normoxia. **A**) V◻O_2_max measured in normoxia and hypoxia. See Fig. 2 for details on hemoglobin (Hb) genotypes and symbols. **B**) There was no correlation between hypoxic V◻O_2_max and arterial O_2_ saturation in hypoxia (P = 0.8103) across individuals (mean ± SEM for each genotype are shown as error bars).

### Hb genotype influenced the acclimation responses of red blood cell *P*_50_ and arterial O_2_ saturation to chronic hypoxia

There were main effects of hypoxia acclimation that tended to increase both red blood cell *P*_50_ (P = 0.0002) and arterial O_2_ saturation measured at V◻O_2_max in hypoxia (P = 0.0005), but the acclimation response appeared to differ between genotypes (Fig. 4, Additional file 3: Table S2). Mice with the lowland α-globin variant exhibited no plasticity in red blood cell *P*_50_ in response to hypoxia acclimation, whereas mice with highland α-globin increased red blood cell *P*_50_ to values that were comparable to mice with lowland α-globin. Conversely, mice with lowland α-globin showed much greater plasticity in arterial O_2_ saturation in hypoxia following hypoxia acclimation, with all individuals increasing saturation (on average by ~13% saturation units). Mice with highland α-globin showed little to no change in saturation after hypoxia acclimation. Hypoxia acclimation increased Hct (P < 0.0001) and [Hb] (P < 0.0001; Fig. 1), but neither these traits nor the Hill coefficient were influenced by Hb genotype (Additional file 4: Fig. S3, Additional file 3: Table S2).

**Figure 4.**
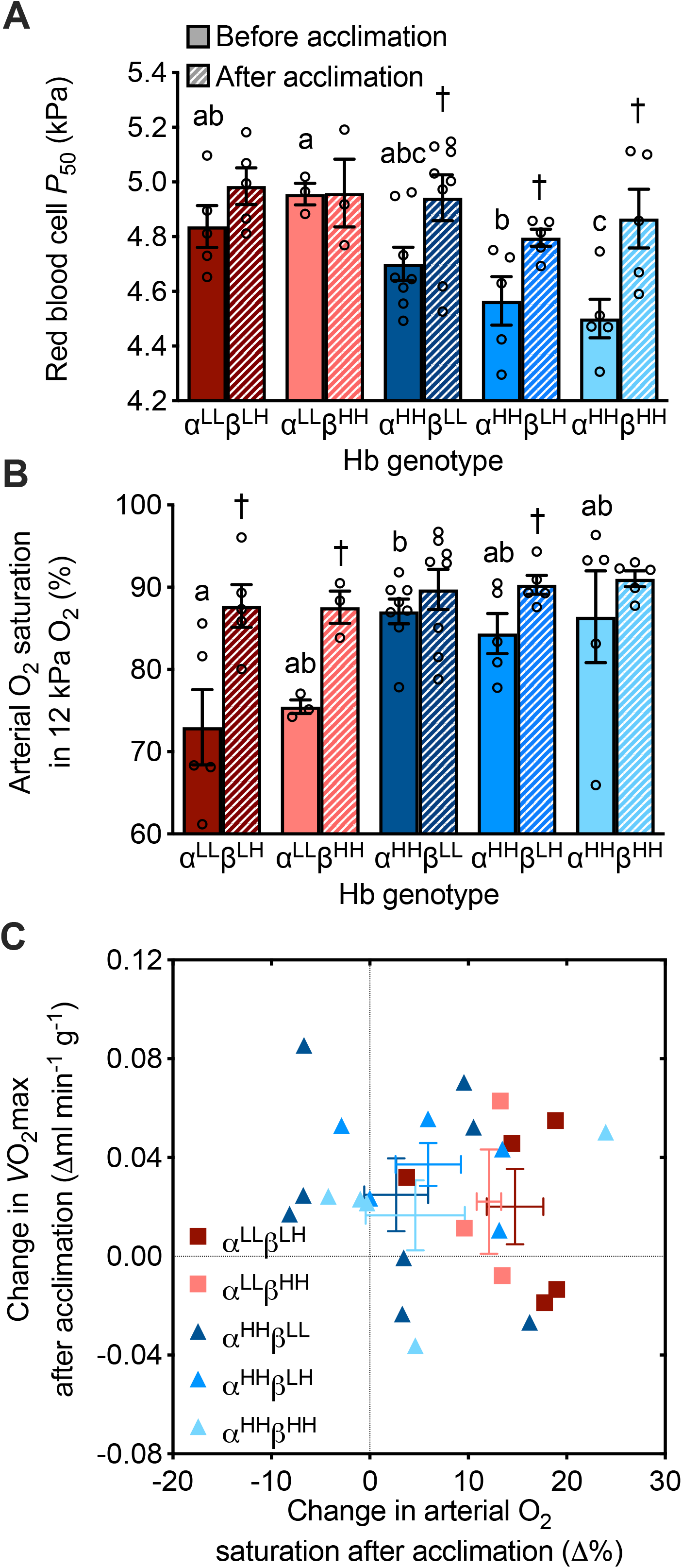
The effects of hypoxia acclimation on red blood cell O_2_ affinity and arterial O_2_ saturation differed between hemoglobin (Hb) genotypes in F_2_ inter-population hybrid deer mice, but the effects of hypoxia acclimation on V◻O_2_max did not. **A**) Red blood cell *P*_50_ (O_2_ pressure at which Hb is 50% saturated) and **B**) arterial O_2_ saturation at V◻O_2_max in hypoxia, measured before and after a 6-wk acclimation to hypobaric hypoxia (12 kPa O_2_). ^†^P < 0.05 vs. pre-acclimation value within a genotype. P < 0.05 between genotypes within an acclimation condition for values not sharing a letter. **C**) The change in hypoxic V◻O_2_max plotted against the change in arterial O_2_ saturation in hypoxia in individuals in response to hypoxia acclimation (mean ± SEM for each genotype are shown as error bars). See Fig. 2 for other details on Hb genotypes.

V◻O_2_max in hypoxia increased after hypoxia acclimation (P = 0.0013), but this response was not influenced by Hb genotype (P = 0.1764; Additional file 6: Fig. S5, Additional file 3: Table S2). The magnitude of change in hypoxic V◻O_2_max following hypoxia acclimation was not associated with the magnitude of change in arterial O_2_ saturation (Fig. 4C). Hypoxia acclimation also increased heart rate (P = 0.0031), total ventilation, (P < 0.0001), tidal volume (P = 0.0005), and breathing frequency (P < 0.0001) measured at V◻O_2_max in hypoxia, but none of these traits were affected by Hb genotype (Additional file 7: Fig. S6, Additional file 3: Table S2). Normoxic V◻O_2_max was not affected by hypoxia acclimation or Hb genotype, nor were the measurements of heart rate, total ventilation, tidal volume, or breathing frequency at normoxic V◻O_2_max affected by Hb genotype (Additional file 8: Fig. S7, Additional file 3: Table S4). However, there was a main effect of genotype on arterial O_2_ saturation measured at V◻O_2_max in normoxia (P = 0.0291) that appeared to result from slightly lower saturation values in mice with characteristic lowland α- and β-globin genotypes (α^LL^β^LH^; Additional file 8: Fig. S7B, Additional file 3: Table S4).

### Sensitivity analysis suggested that effects of Hb-O_2_ affinity on V◻O_2_max in hypoxia are contingent on tissue O_2_ diffusing capacity (D_T_O_2_)

We examined the interactive effects of Hb-O_2_ affinity and D_T_O_2_ on V◻O_2_max in hypoxia using a mathematical model of O_2_ flux through the O_2_ transport pathway. We generated the initial solutions of the model using empirical data collected for deer mice, and then performed a sensitivity analysis to determine the effects of increasing D_T_O_2_ on V◻O_2_max at each of the red blood cell *P*_50_ values for mice with characteristic highland (α^HH^β^HH^) and lowland (α^LL^β^LH^) Hb genotypes. Increasing D_T_O_2_ by 50% increased V◻O_2_max, but the effect was greater with the *P*_50_ of the high-affinity α^HH^β^HH^ genotype (11.8%) than with the lower affinity α^LL^β^LH^ genotype (8.5%; Fig. 5A). The effect of *P*_50_ was accentuated when D_T_O_2_ was increased above 41%, when venous *P*O_2_ (and thus venous O_2_ saturation) fell to zero at the higher *P*_50_ (Fig. 5B). These results indicate that an increase in Hb-O_2_ affinity only contributes to an enhancement of V◻O_2_max in hypoxia if it is paired with an increase in D_T_O_2_ in thermogenic tissues (i.e., skeletal muscle and/or brown adipose tissue).

**Figure 5.**
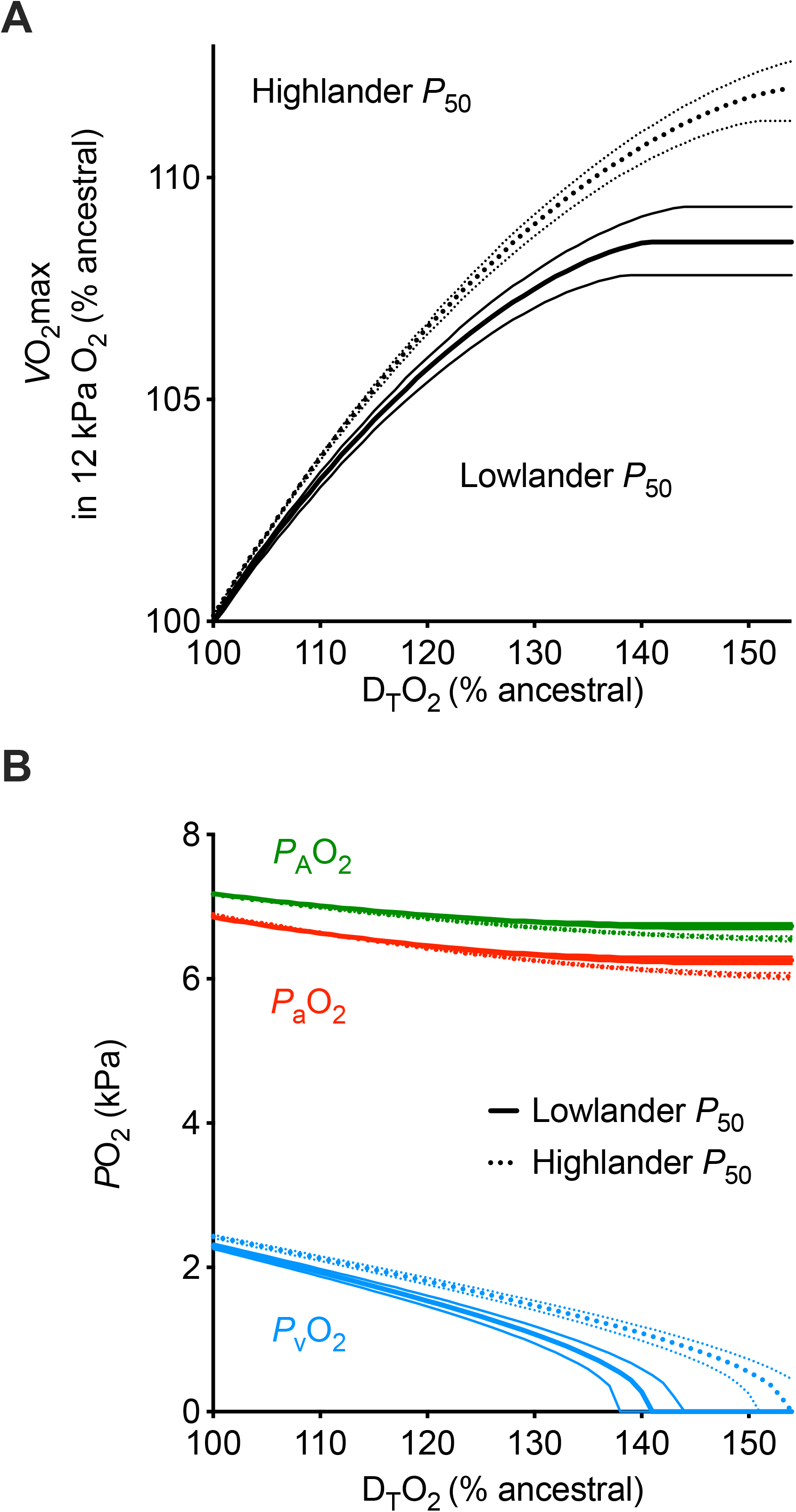
Effects of increasing tissue O_2_ diffusing capacity (D_T_O_2_) on hypoxic V◻O_2_max and O_2_ partial pressures (*P*O_2_) using mathematical modeling of the O_2_ transport pathway. **A**) Relative changes in hypoxic V◻O_2_max and **B**) changes in alveolar (*P*_A_O_2_), arterial (*P*_a_O_2_) and venous (*P*_v_O_2_) *P*O_2_ in response to relative increases in D_T_O_2_. Effects were modeled using the mean (bold lines) ± SEM (fine lines) values of red blood cell *P*_50_ for mice with hemoglobin genotypes that were most characteristic of lowlanders (α^LL^β^LH^) and highlanders (α^HH^β^HH^).

## Discussion

Our study provides evidence that the adaptive benefit of increasing Hb-O_2_ affinity is contingent on the capacity of active tissues to extract O_2_ from the blood. In agreement with previous studies (5, 6, 20, 21, 24, 26, 43), our data from F_2_ inter-population hybrids demonstrate that Hb variants from high-altitude deer mice confer a higher Hb-O_2_ affinity than Hb from lowland conspecifics, and that this evolved increase in affinity augments arterial O_2_ saturation in hypoxia by 9-14%. However, these genetically based changes alone did not augment V◻O_2_max (*i.e.*, aerobic performance) in hypoxia. Modeling of the O_2_ transport pathway revealed that increases in Hb-O_2_ affinity would only be expected to enhance V◻O_2_max in hypoxia if O_2_ diffusing capacity were increased to augment tissue O_2_ extraction. Importantly, recent evidence suggests that high-altitude mice have evolved a highly aerobic skeletal muscle phenotype with an enhanced capacity for O_2_ diffusion (37–41). In particular, the gastrocnemius muscle of highland deer mice has greater capillary density and a redistribution of mitochondria to a subsarcolemmal location that is closer to capillaries, each of which would increase O_2_ diffusing capacity. Our results therefore suggest that increases in both Hb-O_2_ affinity and tissue O_2_ diffusing capacity likely contributed to the adaptive increases in V◻O_2_max in high-altitude deer mice. These findings suggest the testable hypothesis that other hypoxia-adapted, high-altitude vertebrates that have evolved derived increases in Hb-O_2_ affinity will also have evolved increases in tissue capillarity and/or other changes that augment O_2_ diffusing capacity.

The genetically based differences in Hb function led to predictable differences in arterial O_2_ saturation during acute and chronic hypoxia. Amino acid variation in Hb genes is not always associated with changes in O_2_-binding properties (16, 44), and even in cases where it has been possible to document causal effects of specific mutations on Hb function (13, 15, 23–26, 34, 42, 45–48), the *in vivo* effects on blood oxygenation have rarely been examined. Our study suggests that it is critically important to examine how genetic changes in proximal biochemical phenotypes affect higher-level physiological phenotypes (*e.g.*, arterial O_2_ saturation and V◻O_2_max in hypoxia) to fully understand their potential adaptive significance.

Genetic variation in Hb altered the acclimation response to chronic hypoxia, as highland α-globin genotypes were associated with increased plasticity in Hb-O_2_ affinity of red blood cells. This variation was likely a result of differences in sensitivity to 2,3-diphosphoglycerate (2,3-DPG), an allosteric modulator of Hb-O_2_ affinity. Concentrations of 2,3-DPG in erythrocytes are known to increase in response in chronic hypoxia, which tends to reduce red blood cell Hb-O_2_ affinity (49–53). Previous studies have shown that Hb from high-altitude deer mice are more sensitive to 2,3-DPG in the presence of Cl^−^ than Hb from low-altitude mice (24, 26). Therefore, in the current study, if red blood cell concentrations of 2,3-DPG were comparable across genotypes, differences in 2,3-DPG sensitivity could explain the differences in plasticity of red blood cell Hb-O_2_ affinity. This mechanism may also explain why genotypes differed in the magnitude of plasticity in arterial O_2_ saturation in response to chronic hypoxia. Several physiological adjustments contribute to increasing arterial O_2_ saturation after hypoxia acclimation, including increases in total ventilation (Fig. 1) and adjustments in lung function to augment pulmonary O_2_ diffusion (36, 54), and these effects could potentially be counteracted by reductions in red blood cell Hb-O_2_ affinity. Such reductions in affinity did not occur in mice with lowland α-globin, such that they experienced greater improvements in arterial O_2_ saturation after hypoxia acclimation.

Our results indicate that the adaptive benefit of increasing Hb-O_2_ affinity is contingent on the O_2_ diffusing capacity of active tissues. Our study provides empirical evidence that genetically based increases in Hb-O_2_ affinity and arterial O_2_ saturation alone are not sufficient to improve aerobic capacity in hypoxia. We also demonstrate that the adaptive benefit of increasing Hb-O_2_ affinity is contingent on having a tissue O_2_ conductance (D_T_O_2_) that is sufficiently high to take advantage of the greater arterial O_2_ saturation and extract more O_2_ from the blood. The relationship between Hb-O_2_ affinity and V◻O_2_max in hypoxia is a contentious topic (12, 25, 28), with different empirical studies and theoretical models providing contradictory results (5, 27, 29, 30, 33, 55). In fact, previous investigation in deer mice has shown that mice possessing highland α-globin alleles with higher Hb-O_2_ affinity did have higher V◻O_2_max in hypoxia than mice with lowland α-globin haplotypes (5). However, in this previous study (in which genotyping was based on protein electrophoresis), different α-globin alleles were backcrossed into a highland genetic background (5), unlike the current study in which alternative allelic variants were randomized against an admixed highland/lowland background. As discussed above, highland deer mice appear to have evolved a higher capacity for O_2_ diffusion and utilization in skeletal muscles than their lowland conspecifics, comparable to some differences between high-altitude and low-altitude human populations (56). It is therefore possible that the highland mice used in this previous study (5) had a higher D_T_O_2_ than the F_2_ inter-population hybrids used in our present study, which would explain the observed differences in the relative influence of Hb genotype on V◻O_2_max. Indeed, our modeling shows that the adaptive benefits of increasing Hb-O_2_ affinity are critically dependent on D_T_O_2_. Together, our findings suggest adaptive increases in V◻O_2_max in high-altitude deer mice may have been facilitated by evolved increases in D_T_O_2_, which were required in order for increases in Hb-O_2_ affinity to confer an adaptive benefit at high-altitude.

## Conclusions

Complex organismal traits are often the result of multiple interacting genes and phenotypes, but the role of these interactions in shaping adaptive traits is poorly understood. Our findings demonstrate that adaptive increases in thermogenic capacity result from a functional interaction between blood hemoglobin and active tissues, in which the adaptive benefit of increasing hemoglobin O_2_ affinity is contingent on the capacity for O_2_ diffusion from the blood. This helps reconcile controversy about the general role of hemoglobin in hypoxia tolerance, and provides insight into physiological mechanisms of high-altitude adaptation.

## Methods

### Animals

Wild deer mice (*Peromyscus maniculatus*) were live-trapped at high altitude on the summit of Mount Evans (Clear Creed County, CO, USA at 39°35’18”N, 105°38’38”W; 4350 m above sea level) and at low altitude on the Great Plains (Nine Mile Prairie, Lancaster County, NE, USA at 40°52’12”N, 96°48’20.3”W; 430 m above sea level), and were transported to the University of Montana (elevation 978 m). The wild mice were used to produce one family of first-generation inter-population hybrids (F_1_), created by crossing a highland male and a lowland female. These F_1_ hybrids were raised to maturity and used for full-sibling matings to produce 4 families of male and female second-generation hybrid progeny (F_2_). These F_2_ hybrids (n = 26) were raised to adulthood (1-1.5 years old), a small volume of blood was obtained for genotyping (sampled from the facial vein and then stored at −80°C), and mice were then transported to McMaster University (near sea level) for subsequent experiments (see below). Prior to experimentation, all mice were kept in standard holding conditions (24-25°C, 12:12-h light-dark photoperiod) under normal atmospheric conditions, with unlimited access to water and standard mouse chow. All animal protocols were approved by institutional animal research ethics boards.

Each mouse was genotyped for determination of α- and β-globin haplotypes. Tetrameric hemoglobin isoforms of adult *P. maniculatus* incorporate α-chain subunits that are encoded by two tandem gene duplicates, HBA-T1 and HBA-T2 (separated by 5.0 kb on Chromosome 8), and β-chain subunits that are encoded by two other tandem duplicates, HBB-T1 and HBB-T2 (separated by 16.2 kb on Chromosome 1) (34, 57, 58). A reverse-transcriptase PCR (RT-PCR) approach was used to obtain sequence data for all four of the adult-expressed α- and β-globin transcripts (26, 34). Total RNA was extracted from red blood cells using the RNeasy Plus Mini Kit (Qiagen, Valencia, CA, USA). Globin transcripts were then amplified from 1 μg of extracted RNA using the One-Step RT-PCR system with Platinum Taq DNA polymerase High Fidelity (Invitrogen, Carlsbad, CA, USA). PCR cycling was performed with 1 cycle at 50 °C for 30 min, 1 cycle at 95°C for 15 min, 34 cycles at 94°C for 30 s, 55°C for 30 s, and 72°C for 1 min, and then a final extension cycle at 72°C for 3 min. For the α-globin transcripts, the same primer pair was used for HBA-T1 and HBA-T2 (forward: CTGATTCTCACAGACTCAGGAAG, reverse: CCAAGAGGTACAGGTGCGAG). For the β-globin transcripts, the same RT-PCR primer pair was used for HBB-T1 and HBB-T2 (forward: GACTTGCAACCTCAGAAACAGAC, reverse: GACCAAAGGCCTTCATCATTT). Gel-purified RT-PCR products were then cloned into pCR4-TOPO vector using the TOPO TA cloning kit (Invitrogen), and automated DNA sequencing of cloned PCR products was performed using Big Dye chemistry (ABI 3730 capillary sequencer; Applied Biosystems, Foster City, CA, USA). For each mouse, we sequenced 6 clones containing products of HBA-specific RT-PCR, and 6 clones containing products of HBB-specific RT-PCR. Thus, full-length inserts representing cDNAs of all expressed HBA and HBB genes were sequenced at 6-fold coverage, and the haplotype phase of all variable sites was determined experimentally. We thus identified 5 distinct combinations of highland (H)- and lowland (L)-associated α- and β-globin haplotypes: n = 5 α^LL^β^LH^, 2 male, 3 female; n = 3 α^LL^β^HH^, 1 male, 2 female; n = 8 α^HH^β^LL^, 5 male, 3 female; n = 5 α^HH^β^LH^, 5 male; and n = 5 α^HH^β^HH^, 5 male.

### Acclimation treatments

Physiological measurements (see below) were taken for each mouse both before (mean ± SEM body mass, 23.8 ± 0.9 g) and after (23.7 ± 0.8 g) a six-week acclimation to hypobaric hypoxia (approx. 12 kPa *P*O_2_), approximating the O_2_ levels experienced by highland deer mice living at 4,350 m above sea level in the wild (see Additional file 2: Fig. S2 for body masses for each genotype before and after hypoxia acclimation). This was achieved by placing mice into custom-made hypobaric chambers inside which barometric pressure was maintained at 60 kPa using a vacuum pump, as previously described (37, 59). Mice were removed from the chambers for <20 min twice per week for cage cleaning.

### Respirometry, plethysmography, and pulse oximetry

We combined open-flow respirometry, plethysmography and pulse oximetry to simultaneously measure aerobic capacity for thermogenesis (thermogenic V◻O_2_max), pulmonary ventilation, arterial O_2_ saturation, and heart rate during acute cold (−5°C) exposure in heliox (9, 36). We used established methods of measuring thermogenic V◻O_2_max that have been shown to elicit values of V◻O_2_max that equal or exceed those measured during exercise V◻O_2_max in deer mice (22, 60, 61). These measurements were performed twice before mice were acclimated to hypoxia: once in normoxic heliox (21% O_2_, 79% He) and once in hypoxic heliox (12% O_2_, 88% He) in random order. After hypoxia acclimation, both normoxic and hypoxic V◻O_2_max trials were repeated. V◻O_2_max was measured inside a 530-ml plethysmography chamber that has been previously described in detail (62), and was kept in a regulated freezer to maintain the internal chamber temperature at −5°C (measured with a PT-6 thermocouple, Physitemp). Two days prior to initial trials, the neck fur of each mouse was removed with Nair™ hair-removal product to facilitate pulse oximetry.

Immediately before each trial, each mouse was weighed, fitted with a MouseOx pulse oximetry neck collar (Starr Life Sciences, Oakmont, PA, USA), and placed inside the chamber for 10 min of continuous recording. Incurrent gas flowed through the animal chamber at a rate of 1500 ml min^−1^ (controlled by an MFC-2 mass flow controller, Sable Systems, Las Vegas, NV, USA), and was cooled before entering the plethysmograph by passing through copper coils that were also placed in the freezer. Excurrent gas was subsampled at a rate of 200 ml min^−1^, and dried with pre-baked Drierite before passing through O_2_ and CO_2_ analyzers to determine the fractional concentrations of each gas (FoxBox Respirometry System, Sable Systems). Breathing-induced changes in flow across a pneumotachograph in the chamber lid were measured from pressure oscillations in the animal chamber relative to an identical reference chamber using a differential pressure transducer (Validyne DP45; Cancoppas, Mississauga, ON, Canada) and carrier demodulator (Validyne CD15, Cancoppas), and signals were volume-calibrated before each trial with 300-μl injections using a gas-tight syringe. The core body temperature (*T*_b_) of each mouse was obtained (RET-3-ISO; Physitemp, Clifton, NJ, USA) immediately after being removed from the plethysmograph, and then again at room temperature exactly 24 h afterwards, allowing us to estimate *T*_b_ at V◻O_2_max for use in tidal volume calculations (see below) by assuming that *T*_b_ dropped linearly throughout the trial. All data were acquired and recorded using a PowerLab 8/32 and LabChart 8 Pro Software (ADInstruments, Colorado Springs, CO, USA), with the exception of pulse oximetry data, which were obtained using Starr Life Sciences acquisition hardware and software.

Breathing, heart rate, and arterial O_2_ saturation were recorded at thermogenic V◻O_2_max for each trial. O_2_ consumption rate (V◻O_2_) was calculated from gas concentration and flow measurements using established equations (63). V◻O_2_max was defined as the maximal V◻O_2_ measured over a 10-s period during the 10-min trial. V◻O_2_ usually increased to V◻O_2_max within 5-6 min of entering the chamber and would then decline to less than 90% V◻O_2_max, and all mice had depressed *T*_b_ by the end of each 10-min cold exposure. Tidal volume was calculated (in volumes at BTPS) using established equations for the barometric method in flow-through conditions (64, 65). Total ventilation was calculated as the product of tidal volume and breathing frequency.

### Hematology

Hematology was measured both before and after hypoxia acclimation. Blood samples were taken from the facial vein 3 days after V◻O_2_max measurements. We measured Hb content using Drabkin’s reagent (according to instructions from the manufacturer, Sigma-Aldrich) and hematocrit by spinning the blood in capillary tubes at 12,700 *g* for 5 min. The O_2_ affinity of intact erythrocytes was measured using 10 μl blood in 5 ml buffer containing 0.1 M Hepes, 0.05 M EDTA, 0.1 M NaCl, 0.1% bovine serum albumin, and 0.2% antifoaming agent at pH 7.4. Oxygen dissociation curves were generated at 37 °C using a Hemox Analyzer (TCS Scientific), and red blood cell *P*_50_ and Hill coefficient (*n*) were calculated using Hemox Analytic Software.

### Statistics

We used linear mixed effects models to test for the effects of Hb genotype and acclimation condition using the lme4 (66) package in R (v.3.1.3, R Core Team, 2013). We carried out one set of models to examine the fixed effects of Hb genotype in normoxia-acclimated mice, in absence of the effects of hypoxia acclimation, and with inspired *P*O_2_ as an additional fixed effect. We then carried out a second set of models including data from both before and after chronic hypoxia exposure to examine the effects of Hb genotype, hypoxia acclimation, and their interaction. We used a backwards model selection approach, in which initial models included sex, family, and individual subject as random factors, as well as body mass as a covariate. If these terms had P values above 0.1, they were removed by stepwise backward deletion (starting with the term with the highest P value) and the model was re-run until all terms in the model (with the exception of fixed factors and individual subject) had P values below 0.1. Family was thus included in only 6 of the models (see Additional file 3: Tables S1-S4), while the effects of sex were never significant and were removed from all models. Tukey’s HSD *post hoc* tests were performed to test for pairwise differences between genotypes within an acclimation/*P*O_2_ treatment, and between acclimation/*P*O_2_ treatment groups within each genotype. Data are presented as individual values and as mean ± SEM, unless otherwise stated.

### Modeling the O_2_ transport pathway

Mathematical modeling of the O_2_ transport pathway of deer mice was used to determine the interactive effects of blood-O_2_ affinity and tissue O_2_ diffusing capacity on V◻O_2_max in hypoxia. This was done using established equations that have been used previously to build similar models (30, 67–71). The Fick equation describes the diffusion of oxygen from the alveoli into the blood along capillaries in the lung:

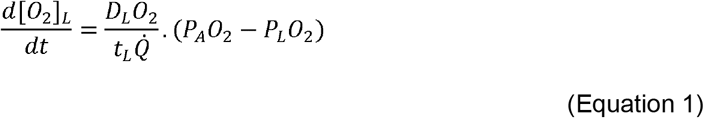

where [O_2_]_L_ and t_L_ are the instantaneous O_2_ content and transit time of the lung capillaries, D_L_O_2_ is the physiological O_2_ diffusing capacity of the lungs, *Q*◻ is cardiac output, and *P*_A_O_2_ and *P*_L_O_2_ are the *P*O_2_ in the alveoli and instantaneously along lung capillaries, respectively. *P*_L_O_2_ began at mixed venous *P*O_2_ (*P*_v_O_2_) and the equation was then integrated over the length on the lung capillaries to determine arterial *P*O_2_ (*P*_a_O_2_) using the Hill equation (Equation 2) to relate [O_2_] and *P*O_2_ in the blood (dependent on blood O_2_ affinity and hemoglobin content):

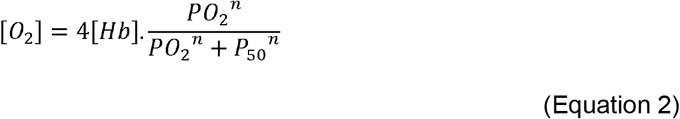

where [Hb] is the hemoglobin content of the blood, *P*_50_ is the *P*O_2_ at which blood is 50% saturated with O_2_, and *n* is the Hill coefficient that describes the cooperativity of blood-O_2_ binding. The Fick equation also describes O_2_ diffusion from the blood in tissue capillaries to the mitochondria:

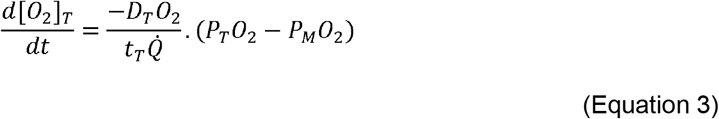

where [O_2_]_T_ and t_T_ are the instantaneous O_2_ content and transit time of the tissue capillaries, D_T_O_2_ is the O_2_ diffusing capacity of the tissues, and *P*_T_O_2_ and *P*_M_O_2_ are the *P*O_2_ instantaneously along the tissue capillaries and at mitochondria, respectively. *P*_M_O_2_ was set to zero to facilitate modeling, but mitochondrial *P*O_2_ is likely quite low and relatively close to zero at V◻O_2_max (69). In this case, *P*_T_O_2_ begins at *P*_a_O_2_ and the equation is integrated along the tissue capillaries to determine *P*_v_O_2_. Mass conservation then matches V◻O_2_max measured from O_2_ extraction at the lungs to that at tissues:

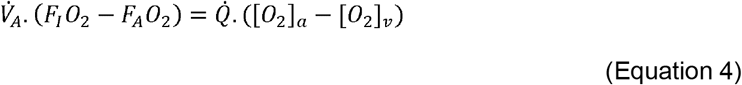

where V◻_A_ is alveolar ventilation, F_I_O_2_ and F_A_O_2_ and the O_2_ fractions of inspired and alveolar gas, respectively, and [O_2_]_a_ and [O_2_]_v_ are arterial and venous oxygen content, respectively.

The above equations were solved using an iterative approach for the key unknown outcome variables, *P*_A_O_2_, *P*_a_O_2_, and *P*_v_O_2_, from which V◻O_2_max was calculated. This was achieved using a F_I_O_2_ of 0.123, our empirical measurements of total ventilation at hypoxic V◻O_2_max, red blood cell *P*_50_ and *n*, and [Hb] in normoxia acclimated α^LL^β^LH^ mice, a body mass specific lung dead space volume of 6.4 μl g^−1^ (72) to calculate alveolar ventilation from total ventilation, and cardiac output at hypoxic V◻O_2_max from our previous measurements in low-altitude deer mice (36). Values for D_L_O_2_ and D_T_O_2_ were chosen by trial and error to re-produce *in vivo* measurements of *P*_a_O_2_, *P*_v_O_2_, and V◻O_2_max. All of the initial values used to solve the model are listed in Additional file 3: Table S5. The model was solved iteratively as follows. Using previously recorded *in vivo P*_v_O_2_ (36) and an initial estimate of *P*_A_O_2_ as a starting point, Equation 1 was integrated to calculate a predicted value of *P*_a_O_2_. This *P*_a_O_2_ value was then used to calculate *P*_v_O_2_ by integrating Equation 3. The two above steps were repeated until the *P*_a_O_2_ and *P*_v_O_2_ values became stable to within 0.05% (< 10 iterations). V◻O_2_max was then calculated using both the left and right sides of Equation 4. If the values did not agree to within 0.05%, *P*_A_O_2_ was adjusted and the above steps were repeated until the left and right sides of Equation 4 were equal to within 0.05%. Reaching a stable solution of the model generally took less than 10 iterations, and the final outcome was independent of the starting estimate of *P*_v_O_2_.

We conducted a sensitivity analysis of the effects of increasing D_T_O_2_ on V◻O_2_max using the mean for the most ancestral ‘lowland’ *P*_50_ that was measured in α^LL^β^LH^ mice, and then again using the mean for the ‘highland’ *P*_50_ that was measured in α^HH^β^HH^ mice, with all other parameters in the model kept constant (including the potential effects of variation in blood pH on *P*_50_), with the exception of *P*_A_O_2_, *P*_a_O_2_, and *P*_v_O_2_ (which were under the influence of the changes in D_T_O_2_ and *P*_50_). We also used the mean + SEM and mean - SEM values of *P*_50_ for each genotype in order to examine the influence of biological variation in *P*_50_ within each genotype. V◻O_2_max and D_T_O_2_ values are expressed here relative to the initial solution generated using the data here from α^LL^β^LH^ mice or from previous measurements in lowland deer mice (Additional file 3: Table S5), which we have termed the ‘ancestral’ values. The calculations were carried out using spreadsheet software (Microsoft Excel), as in some previous models of the O_2_ transport pathway (71), and we have included the spreadsheet template with values for the initial lowland *P*_50_ model (Additional file 9: Dataset and Modeling).

## Supporting information

Additional file 9: Dataset and modeling

## Declarations

### Ethics approval and consent to participate.

All animal protocols were approved by institutional animal research ethics boards.

### Consent for publication.

Not applicable

### Availability of data and materials.

The dataset supporting the conclusions of this article is included within the article and its additional files.

### Author’s contributions

GRS, JFS, and ZAC conceived and designed the study. OHW, CMI, JPV, NG-P, SCC-S, and CN acquired the data. OHW analyzed the data. OHW and GRS interpreted the data and drafted the manuscript. All authors read and approved the final manuscript.

## Acknowledgements

Not applicable.

## Legends to Combined Additional files

**Additional file 1: Figure S1.** Graphical overview of the experimental design of our study. Deer mice from high-(H) and low-(L) altitude populations were crossed in captivity to produce F_1_ interpopulation hybrids that were then mated with siblings to produce the F_2_ interpopulation hybrids that were used in our experiments before and after a 6-wk acclimation to hypobaric hypoxia (12 kPa O_2_). These hybrids were grouped based on the altitudinal origin of their α- and β-globin genotype.

**Additional file 2: Figure S2.** Body mass of F_2_ inter-population hybrids both before and after a 6-wk acclimation to hypobaric hypoxia. Each individual’s mass was measured before normoxic and hypoxic V◻O_2_max trials, with the mean of these values used to create each individual’s data point in the figure. Different α- and β-globin genotypes are shown as superscripts with ^‘L’^ representing the lowland haplotype and ^‘H’^ representing the highland haplotype. There was no effect of genotype (P = 0.2977), acclimation (P = 0.4018), or their interaction (P = 0.3362) on body mass. Bars display mean ± SEM (n = 3-8) with individual data superimposed (circles).

**Additional file 3: Tables S1-S5.** Table S1 - Effects of inspired *P*O_2_ and acclimation to hypoxia on cardiorespiratory physiology of F_2_ inter-population hybrid deer mice at V◻O_2_max, without accounting for effects of genotype. Table S2 - Effects of acclimation to hypoxia and globin genotype on cardiorespiratory physiology in hypoxia of F_2_ inter-population hybrid deer mice. Table S3 - Effects of inspired *P*O_2_ and hemoglobin genotype on cardiorespiratory physiology of F_2_ inter-population hybrid deer mice acclimated to normoxia. Table S4 - Effects of acclimation to hypoxia and globin genotype on cardiorespiratory physiology in normoxia of F_2_ inter-population hybrid deer mice. Table S5 - Parameters used to generate the initial solution in the model of the oxygen transport pathway representing the ‘ancestral condition’ with the most lowland *P*_50_.

**Additional file 4: Figure S3.** Hematology of F_2_ inter-population hybrids measured before and after a 6-wk acclimation to hypobaric hypoxia (12 kPa O_2_). Hct, hematocrit; [Hb], blood hemoglobin content. Different α- and β-globin genotypes are shown as superscripts with ‘^L^’ representing the lowland haplotype and ‘^H^’ representing the highland haplotype. ^†^P < 0.05 vs. pre-acclimation value within a genotype. Bars display mean ± SEM (n = 3-8) with individual data superimposed (circles).

**Additional file 5: Figure S4.** Physiological parameters for F_2_ inter-population hybrids acclimated to normoxia, measured at V◻O_2_max in normoxia (21 kPa O_2_) and hypoxia (12 kPa O_2_). Different α- and β-globin genotypes are shown as superscripts with ‘^L^’ representing the lowland haplotype and ‘^H^’ representing the highland haplotype. *P < 0.05 vs. normoxia value within a genotype. P < 0.05 between genotypes for hypoxic values not sharing a letter. Bars display mean ± SEM (n = 3-8) with individual data superimposed (circles).

**Additional file 6: Figure S5.** Hypoxic V◻O_2_max before and after a 6-wk acclimation to hypobaric hypoxia (12 kPa O_2_). Different α- and β-globin genotypes are shown as superscripts with ‘^L^’ representing the lowland haplotype and ‘^H^’ representing the highland haplotype. ^†^P < 0.05 vs. pre-acclimation value within a genotype. Bars display mean ± SEM (n = 3-8) with individual data superimposed (circles).

**Additional file 7: Figure S6.** Physiological parameters for F_2_ inter-population hybrids measured at V◻O_2_max in hypoxia (12 kPa O_2_) both before and after a 6-wk acclimation to hypobaric hypoxia. Different α- and β-globin genotypes are shown as superscripts with ‘^L^’ representing the lowland haplotype and ‘^H^’ representing the highland haplotype. ^†^P < 0.05 vs. pre-acclimation value within a genotype. P < 0.05 between genotypes within an acclimation condition for values not sharing a letter. Bars display mean ± SEM (n = 3-8) with individual data superimposed (circles).

**Additional file 8: Figure S7.** Physiological parameters for F_2_ inter-population hybrids measured at V◻O_2_max in normoxia (21 kPa O_2_) both before and after a 6-wk acclimation to hypobaric hypoxia. Different α- and β-globin genotypes are shown as superscripts with ‘^L^’ representing the lowland haplotype and ‘^H^’ representing the highland haplotype. ^†^P < 0.05 vs. pre-acclimation value within a genotype. P < 0.05 between genotypes within an acclimation condition for values not sharing a letter. Bars display mean ± SEM (n = 3-8) with individual data superimposed (circles).

**Additional file 9: Dataset and modeling.**

## References

1. Gould SJ. The Structure of Evolutionary Theory. Cambridge, MA, USA.: Harvard Univ. Press; 2002.

2. Jernigan RW, Culver DC, Fong DW. The dual role of selection and evolutionary history as reflected in genetic correlations. Evolution. 1994;48(3):587–96.

3. Wagner GP, Altenberg L. Complex adaptations and the evolution of evolvability. Evolution. 1996;50(3):967–76.

4. Hayes JP, O’Connor CS. Natural selection on thermogenic capacity of high-altitude deer mice. Evolution. 1999;53(4):1280–7.

5. Chappell MA, Snyder LRG. Biochemical and physiological correlates of deer mouse alpha-chain hemoglobin polymorphisms. Proc Natl Acad Sci USA. 1984;81(17):5484–8.

6. Chappell MA, Hayes JP, Snyder LRG. Hemoglobin polymorphisms in deer mice (Peromyscus maniculatus): physiology of beta-globin variants and alpha-globin recombinants. Evolution. 1988;42(4):681–8.

7. Cheviron ZA, Bachman GC, Connaty AD, McClelland GB, Storz JF. Regulatory changes contribute to the adaptive enhancement of thermogenic capacity in high-altitude deer mice. Proc Natl Acad Sci USA. 2012;109(22):8635–40.

8. Cheviron ZA, Bachman GC, Storz JF. Contributions of phenotypic plasticity to differences in thermogenic performance between highland and lowland deer mice. J Exp Biol. 2013;216(7):1160–6.

9. Tate KB, Ivy CM, Velotta JP, Storz JF, McClelland GB, Cheviron ZA, et al. Circulatory mechanisms underlying adaptive increases in thermogenic capacity in high-altitude deer mice. J Exp Biol. 2017;220(20):3616–20.

10. Ivy CM, Scott GR. Control of breathing and the circulation in high-altitude mammals and birds. Comp Biochem Physiol, Part A Mol Integr Physiol. 2015;186:66–74.

11. Storz JF, Scott GR, Cheviron ZA. Phenotypic plasticity and genetic adaptation to high-altitude hypoxia in vertebrates. J Exp Biol. 2010;213(24):4125–36.

12. Storz JF. Hemoglobin–oxygen affinity in high-altitude vertebrates: is there evidence for an adaptive trend? J Exp Biol. 2016;219(20):3190–203.

13. Weber RE. High-altitude adaptations in vertebrate hemoglobins. Respir Physiol Neurobiol. 2007;158(2):132–42.

14. Storz JF. Hemoglobin: insights into protein structure, function, and evolution. Oxford, UK: Oxford University Press; 2019.

15. Natarajan C, Inoguchi N, Weber RE, Fago A, Moriyama H, Storz JF. Epistasis among adaptive mutations in deer mouse hemoglobin. Science. 2013;340(6138):1324–7.

16. Natarajan C, Projecto-Garcia J, Moriyama H, Weber RE, Muñoz-Fuentes V, Green AJ, et al. Convergent Evolution of Hemoglobin Function in High-Altitude Andean Waterfowl Involves Limited Parallelism at the Molecular Sequence Level. PLoS Genet. 2015;11(12):e1005681–e.

17. Natarajan C, Hoffmann FG, Weber RE, Fago A, Witt CC, Storz JF. Predictable convergence in hemoglobin function has unpredictable molecular underpinnings. Science. 2016;354(6310):336–9.

18. Natarajan C, Jendroszek A, Kumar A, Weber RE, Tame JRH, Fago A, et al. Molecular basis of hemoglobin adaptation in the high-flying bar-headed goose. PLoS Genet. 2018;14(4):e1007331.

19. McClelland GB, Scott GR. Evolved mechanisms of aerobic performance and hypoxia resistance in high-altitude natives. Annu Rev Physiol. 2019;81(1):561–83.

20. Snyder LRG. Deer mouse hemoglobins: is there genetic adaptation to high altitude? BioScience. 1981;31(4):299–304.

21. Snyder LRG, Born S, Lechner AJ. Blood oxygen affinity in high- and low-altitude populations of the deer mouse. Respir Physiol. 1982;48(1):89–105.

22. Storz JF, Cheviron ZA, McClelland GB, Scott GR. Evolution of physiological performance capacities and environmental adaptation: insights from high-elevation deer mice (Peromyscus maniculatus). J Mammal. 2019;100(3):910–22.

23. Storz JF, Natarajan C, Cheviron ZA, Hoffmann FG, Kelly JK. Altitudinal variation at duplicated β-globin genes in deer mice: effects of selection, recombination, and gene conversion. Genetics. 2012;190(1):203–16.

24. Storz JF, Runck AM, Sabatino SJ, Kelly JK, Ferrand N, Moriyama H, et al. Evolutionary and functional insights into the mechanism underlying high-altitude adaptation of deer mouse hemoglobin. Proc Natl Acad Sci USA. 2009;106(34):14450–5.

25. Winslow RM. The role of hemoglobin oxygen affinity in oxygen transport at high altitude. Respir Physiol Neurobiol. 2007;158(2):121–7.

26. Storz JF, Runck AM, Moriyama H, Weber RE, Fago A. Genetic differences in hemoglobin function between highland and lowland deer mice. J Exp Biol. 2010;213(15):2565–74.

27. Brauner CJ, Wang T. The optimal oxygen equilibrium curve: a comparison between environmental hypoxia and anemia. Am Zool. 2015;37(1):101–8.

28. Dempsey JA. With haemoglobin as with politics – should we shift right or left? J Physiol. 2020;598(8):1419–20.

29. Wang T, Malte H. O2 uptake and transport: the optimal P50. In: Farrell AP, editor. Encyclopedia of Fish Physiology: from Genome to Environment. Amsterdam: Elsevier; 2011. p. 1845–55.

30. Wagner PD. Insensitivity of VJO2max to hemoglobin-P50 at sea level and altitude. Respir Physiol. 1997;107(3):205–12.

31. Bunn HF. Regulation of hemoglobin function in mammals. Am Zool. 1980;20(1):199–211.

32. Hebbel RP, Eaton JW, Kronenberg RS, Zanjani ED, Moore LG, Berger EM. Human llamas - adaptation to altitude in subjects with high hemoglobin oxygen-affinity. J Clin Invest. 1978;62(3):593–600.

33. Dominelli PB, Wiggins CC, Baker SE, Shepherd JRA, Roberts SK, Roy TK, et al. Influence of high affinity haemoglobin on the response to normoxic and hypoxic exercise. J Physiol. 2020;598(8):1475–90.

34. Natarajan C, Hoffmann FG, Lanier HC, Wolf CJ, Cheviron ZA, Spangler ML, et al. Intraspecific polymorphism, interspecific divergence, and the origins of function-altering mutations in deer mouse hemoglobin. Mol Biol Evol. 2015;32(4):978–97.

35. Bedford NL, Hoekstra HE. Peromyscus mice as a model for studying natural variation. Elife. 2015;4:e06813.

36. Tate KB, Wearing OH, Ivy CM, Cheviron ZA, Storz JF, McClelland GB, et al. Coordinated changes across the O2 transport pathway underlie adaptive increases in thermogenic capacity in high-altitude deer mice. Proc R Soc London, B, Biol Sci. 2020;287(1927):20192750.

37. Lui MA, Mahalingam S, Patel P, Connaty AD, Ivy CM, Cheviron ZA, et al. High-altitude ancestry and hypoxia acclimation have distinct effects on exercise capacity and muscle phenotype in deer mice. Am J Physiol Regul Integr Comp Physiol. 2015;308(9):R779–R91.

38. Scott GR, Elogio TS, Lui MA, Storz JF, Cheviron ZA. Adaptive modifications of muscle phenotype in high-altitude deer mice are associated with evolved changes in gene regulation. Mol Biol Evol. 2015;32(8):1962–76.

39. Dawson NJ, Lyons SA, Henry DA, Scott GR. Effects of chronic hypoxia on diaphragm function in deer mice native to high altitude. Acta Physiol. 2018;223(1):16.

40. Mahalingam S, McClelland GB, Scott GR. Evolved changes in the intracellular distribution and physiology of muscle mitochondria in high-altitude native deer mice. J Physiol. 2017;595(14):4785–801.

41. Scott GR, Guo KH, Dawson NJ. The mitochondrial basis for adaptive variation in aerobic performance in high-altitude deer mice. Integr Comp Biol. 2018;58(3):506–18.

42. Storz JF, Sabatino SJ, Hoffmann FG, Gering EJ, Moriyama H, Ferrand N, et al. The molecular basis of high-altitude adaptation in deer mice. PLoS Genet. 2007;3(3):e45–e.

43. Jensen B, Storz JF, Fago A. Bohr effect and temperature sensitivity of hemoglobins from highland and lowland deer mice. Comp Biochem Physiol, Part A Mol Integr Physiol. 2016;195:10–4.

44. Cheviron ZA, Natarajan C, Projecto-Garcia J, Eddy DK, Jones J, Carling MD, et al. Integrating Evolutionary and Functional Tests of Adaptive Hypotheses: A Case Study of Altitudinal Differentiation in Hemoglobin Function in an Andean Sparrow, Zonotrichia capensis. Mol Biol Evol. 2014;31(11):2948–62.

45. Poyart C, Wajcman H, Kister J. Molecular adaptation of hemoglobin function in mammals. Respir Physiol. 1992;90(1):3–17.

46. Storz JF. Hemoglobin function and physiological adaptation to hypoxia in high-altitude mammals. J Mammal. 2007;88(1):24–31.

47. Storz JF, Moriyama H. Mechanisms of hemoglobin adaptation to high altitude hypoxia. High Alt Med Biol. 2008;9(2):148–57.

48. Weber RE, Fago A. Functional adaptation and its molecular basis in vertebrate hemoglobins, neuroglobins and cytoglobins. Respir Physiol Neurobiol. 2004;144(2):141–59.

49. Lenfant C, Torrance J, English E, Finch CA, Reynafarje C, Ramos J, et al. Effect of altitude on oxygen binding by hemoglobin and on organic phosphate levels. J Clin Invest. 1968;47(12):2652–6.

50. Lenfant C, Torrance JD, Reynafarje C. Shift of the O2-Hb dissociation curve at altitude: mechanism and effect. J Appl Physiol. 1971;30(5):625–31.

51. Milles JJ, Chesner IM, Oldfield S, Bradwell AR. Effect of acetazolamide on blood gases and 2,3 DPG during ascent and acclimatization to high altitude. Postgrad Med J. 1987;63(737):183–4.

52. Mairbaurl H, Oelz O, Bartsch P. Interactions between Hb, Mg, DPG, ATP, and Cl determine the change in Hb-O2 affinity at high altitude. J Appl Physiol. 1993;74(1):40–8.

53. Savourey G, Launay J-C, Besnard Y, Guinet A, Bourrilhon C, Cabane D, et al. Control of erythropoiesis after high altitude acclimatization. Eur J Appl Physiol. 2004;93(1):47–56.

54. Ivy CM, Scott GR. Control of breathing and ventilatory acclimatization to hypoxia in deer mice native to high altitudes. Acta Physiol. 2017;221(4):266–82.

55. Woodson RD, Auerbach S. Effect of increased oxygen affinity and anemia on cardiac output and its distribution. J Appl Physiol. 1982;53(5):1299–306.

56. Gilbert-Kawai E, Coppel J, Court J, Kaaij Jvd, Vercueil A, Feelisch M, et al. Sublingual microcirculatory blood flow and vessel density in Sherpas at high altitude. J Appl Physiol. 2017;122(4):1011–8.

57. Hoffmann FG, Opazo JC, Storz JF. New Genes Originated via Multiple Recombinational Pathways in the β-Globin Gene Family of Rodents. Mol Biol Evol. 2008;25(12):2589–600.

58. Storz JF, Hoffmann FG, Opazo JC, Moriyama H. Adaptive Functional Divergence Among Triplicated α-Globin Genes in Rodents. Genetics. 2008;178(3):1623–38.

59. McClelland GB, Hochachka PW, Weber J-M. Carbohydrate utilization during exercise after high-altitude acclimation: A new perspective. Proc Natl Acad Sci USA. 1998;95(17):10288–93.

60. Chappell MA, Hammond KA. Maximal aerobic performance of deer mice in combined cold and exercise challenges. J Comp Physiol B. 2004;174(1):41–8.

61. Rosenmann M, Morrison P. Maximum oxygen consumption and heat loss facilitation in small homeotherms by He-O2. Am J Physiol. 1974;226(3):490–5.

62. Ivy CM, Scott GR. Ventilatory acclimatization to hypoxia in mice: methodological considerations. Respir Physiol Neurobiol. 2017;235:95–103.

63. Lighton JRB. Measuring metabolic rates: a manual for scientists: OUP Oxford; 2018.

64. Drorbaug JE, Fenn WO. A barometric method for measuring ventilation in newborn infants. Pediatrics. 1955;16(1):81–7.

65. Jacky JP. Barometric measurement of tidal volume: effects of pattern and nasal temperature. J Appl Physiol. 1980;49(2):319–25.

66. Bates D, Machler M, Bolker BM, Walker SC. Fitting linear mixed-effects models using lme4. J Stat Softw. 2015;67(1):1–48.

67. Wagner PD. Algebraic analysis of the determinants of VJO2,max. Respir Physiol. 1993;93(2):221–37.

68. Wagner PD. Determinants of maximal oxygen transport and utilization. Annu Rev Physiol. 1996;58(1):21–50.

69. Wagner PD. A theoretical analysis of factors determining VJO2max at sea level and altitude. Respir Physiol. 1996;106(3):329–43.

70. Scott GR, Milsom WK. Flying high: a theoretical analysis of the factors limiting exercise performance in birds at altitude. Respir Physiol Neurobiol. 2006;154(1):284–301.

71. Wang T, Hicks JW. An integrative model to predict maximum O2 uptake in animals with central vascular shunts. Zoology. 2002;105(1):45–53.

72. Fallica J, Das S, Horton M, Mitzner W. Application of carbon monoxide diffusing capacity in the mouse lung. J Appl Physiol. 2011;110(5):1455–9.

