## Supplementary figures and images for "The adaptive benefit of evolved increases in hemoglobin-O_2_ affinity is contingent on tissue O_2_ diffusing capacity in high-altitude deer mice"

### Additional file 1: Figure S1

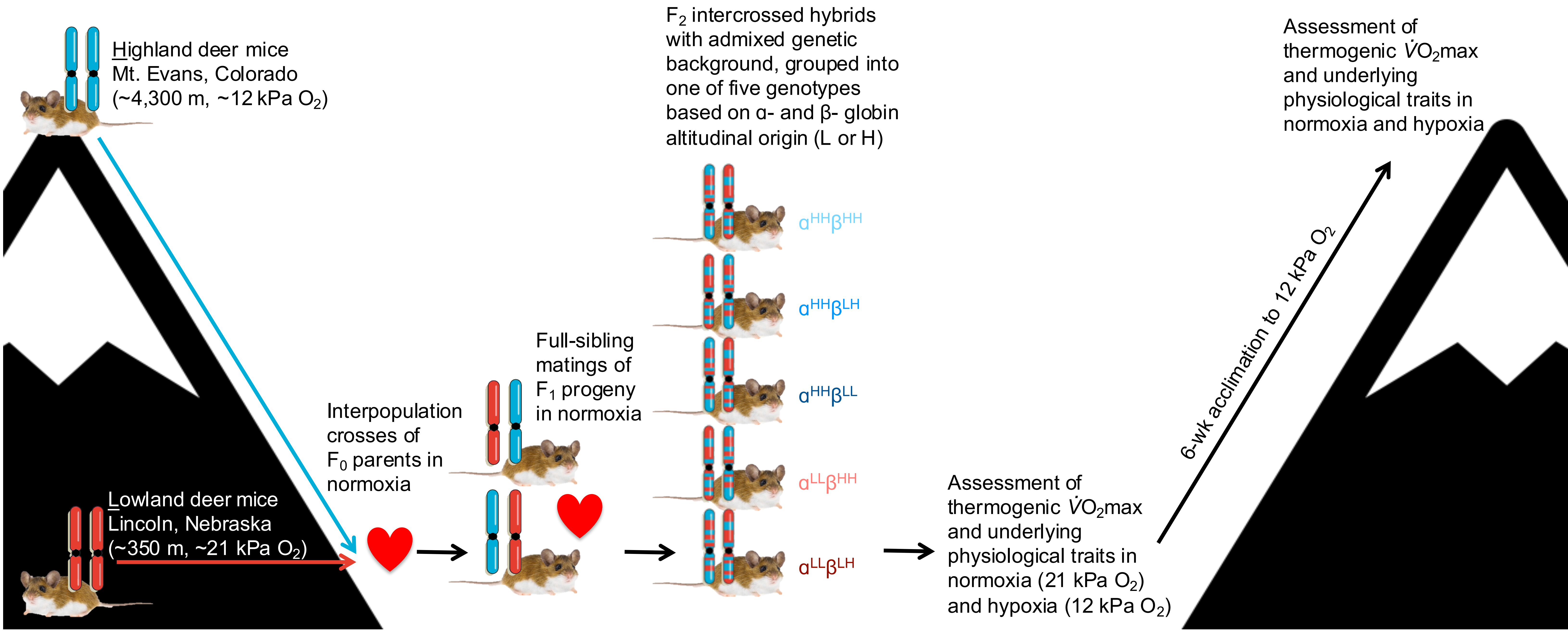

### Additional file 2: Figure S2

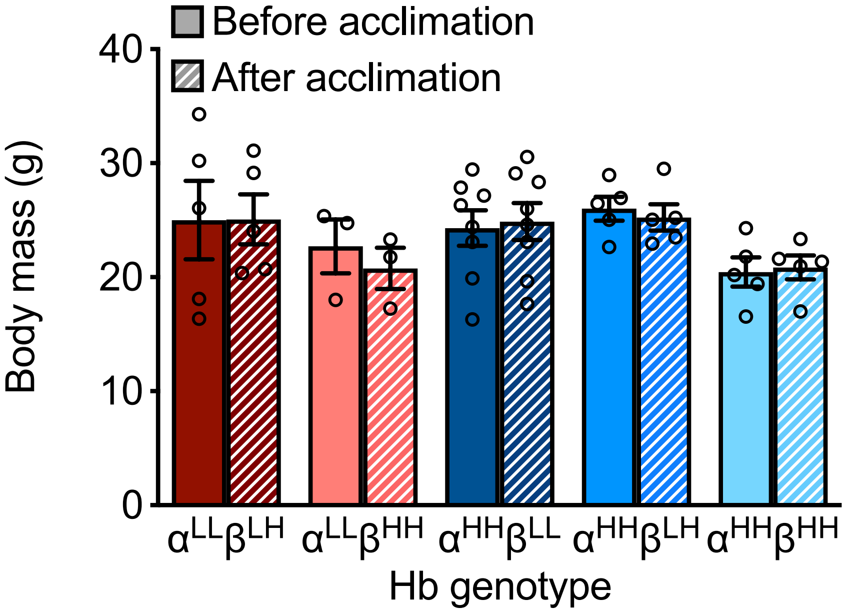

### Additional file 4: Figure S3

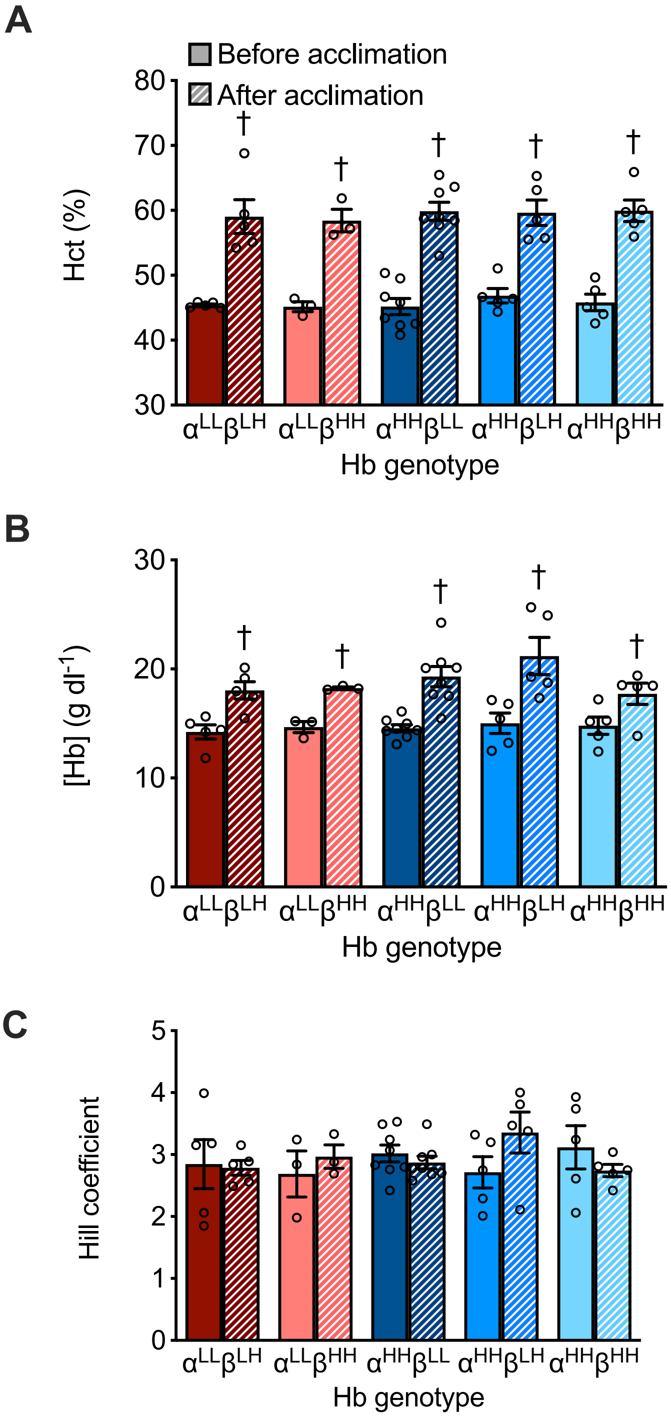

### Additional file 5: Figure S4

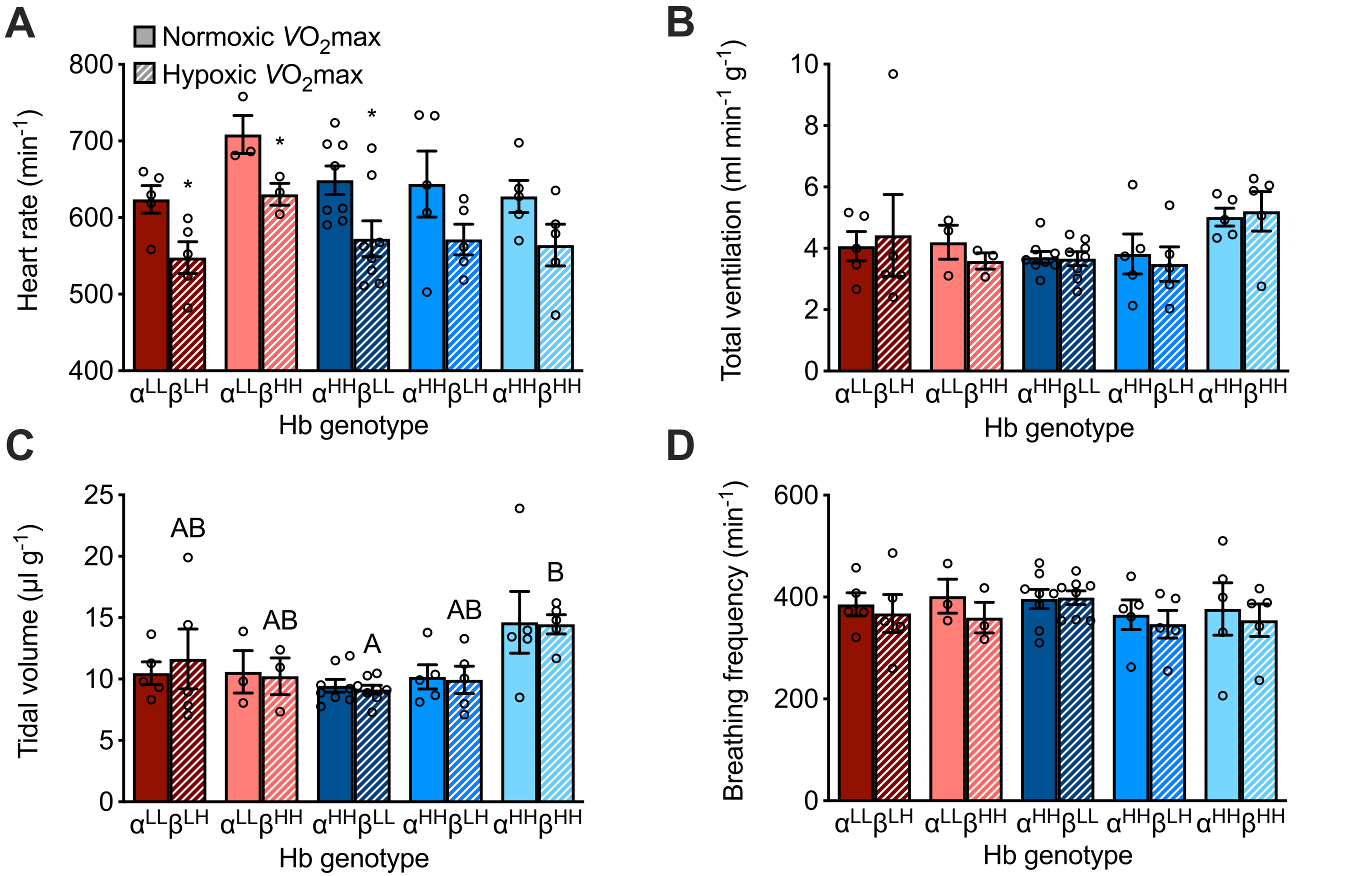

### Additional file 6: Figure S5

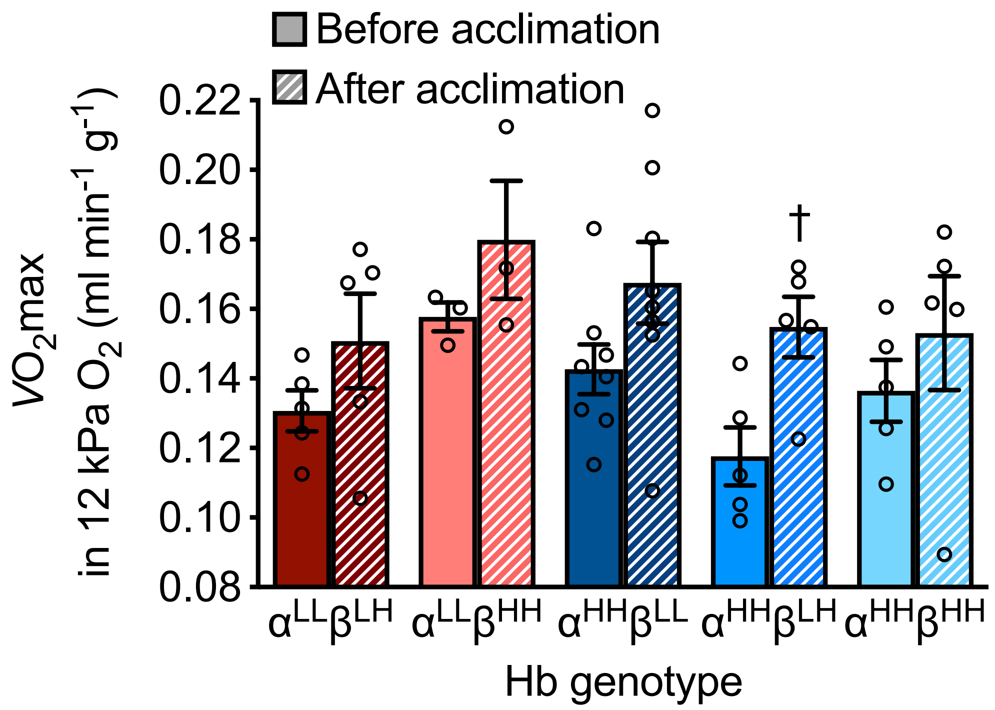

### Additional file 7: Figure S6

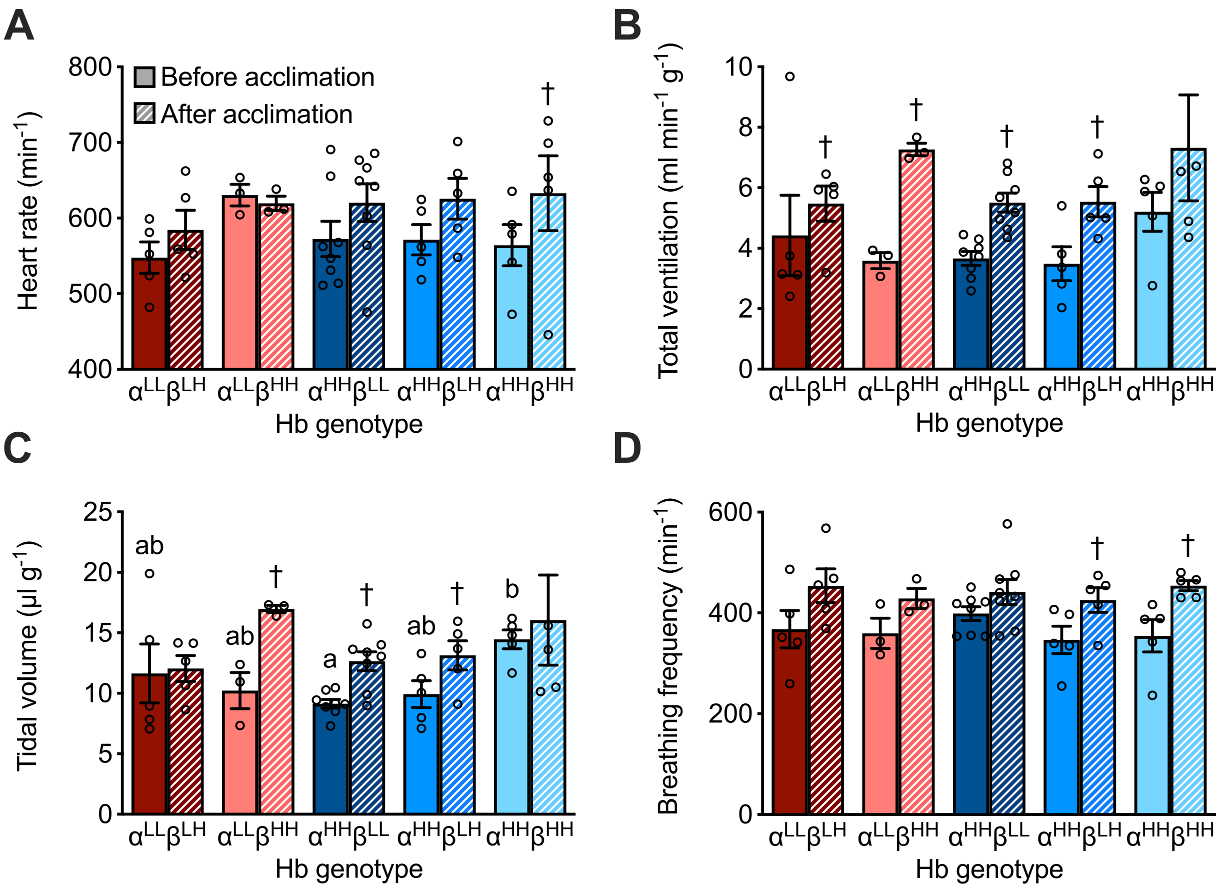

### Additional file 8: Figure S7

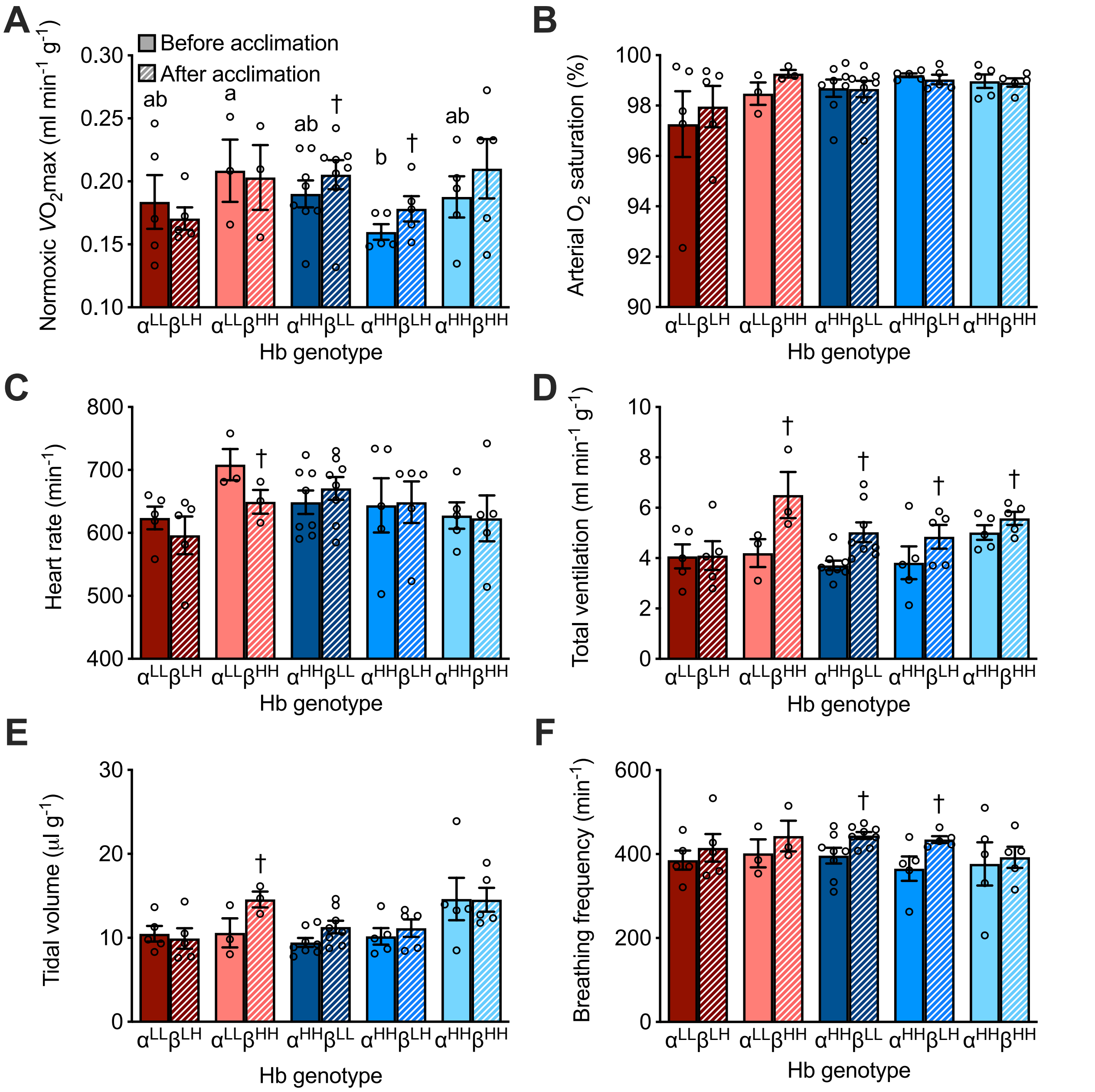
