## Additional file 3: Tables S1-S5 for "The adaptive benefit of evolved increases in hemoglobin-O_2_ affinity is contingent on tissue O_2_ diffusing capacity in high-altitude deer mice"

**Table S1.** Effects of inspired $PO_2$ and acclimation to hypoxia on cardiorespiratory physiology of $F_2$ inter-population hybrid deer mice at $\dot{V}O_2max$, without accounting for effects of genotype.

| Trait | Animal mass | Acclimation | $PO_2$ | Interaction[1] |
|---|---|---|---|---|
| $\dot{V}O_2max$[2] | $F_{1,98} = 37.9184$<br>$P < 0.0001$* | $F_{1,98} = 16.8561$<br>$P = 0.0001$* | $F_{1,98} = 94.4737$<br>$P < 0.0001$* | NS |
| Arterial $O_2$ saturation | $F_{1,98} = 3.0602$<br>$P = 0.0909$ | $F_{1,98} = 14.5282$<br>$P = 0.0003$* | $F_{1,98} = 181.5338$<br>$P < 0.0001$* | $F_{1,98} = 13.0275$<br>$P = 0.0006$* |
| Heart rate[2] | $F_{1,97} = 4.3086$<br>$P = 0.0457$* | $F_{1,97} = 4.6517$<br>$P = 0.0342$* | $F_{1,97} = 27.5056$<br>$P < 0.0001$* | $F_{1,97} = 7.6175$<br>$P = 0.0073$* |
| Total ventilation | NS | $F_{1,99} = 49.9311$<br>$P < 0.0001$* | $F_{1,99} = 3.5126$<br>$P = 0.0648$ | $F_{1,99} = 5.5837$<br>$P = 0.0207$* |
| Tidal volume | NS | $F_{1,99} = 22.2705$<br>$P < 0.0001$* | $F_{1,99} = 3.3406$<br>$P = 0.0716$ | $F_{1,99} = 5.1393$<br>$P = 0.0263$* |
| Breathing frequency | NS | $F_{1,100} = 24.9573$<br>$P < 0.0001$* | $F_{1,100} < 0.0001$<br>$P = 0.9936$ | NS |

*$P < 0.05$. [1]Interaction between acclimation and inspired $PO_2$. NS denotes no significant effect of the factor, which was removed from the final statistical model. [2]Mouse family was also included as a random factor in the mixed model as $0.05 < P < 0.1$.

**Table S2.** Effects of acclimation to hypoxia and globin genotype on cardiorespiratory physiology in hypoxia of $F_2$ inter-population hybrid deer mice.

| Trait | Animal mass | Acclimation | Hb genotype | Interaction[1] |
|---|---|---|---|---|
| Hypoxic $\dot{V}O_2$max | $F_{1,41}$ = 28.0073<br>P < 0.0001* | $F_{1,41}$ = 13.0967<br>P = 0.0013* | $F_{4,41}$ = 1.7576<br>P =0.1764 | NS |
| Arterial $O_2$ saturation | NS | $F_{1,42}$ = 15.8951<br>P = 0.0005* | $F_{4,42}$ = 3.0259<br>P = 0.0407* | NS |
| Heart rate[2] | NS | $F_{1,41}$ = 10.7106<br>P = 0.0031* | $F_{4,41}$ = 0.7020<br>P = 0.5999 | NS |
| Total ventilation | NS | $F_{1,42}$ = 34.0654<br>P < 0.0001* | $F_{4,42}$ = 0.3396<br>P = 0.8481 | NS |
| Tidal volume | NS | $F_{1,42}$ = 15.9684<br>P = 0.0005* | $F_{4,42}$ = 0.6548<br>P = 0.6300 | NS |
| Breathing frequency | NS | $F_{1,42}$ = 23.7780<br>P < 0.0001* | $F_{4,42}$ = 0.5053<br>P = 0.7323 | NS |
| Hematocrit | NS | $F_{1,42}$ = 222.7637<br>P < 0.0001* | $F_{4,42}$ = 0.2031<br>P = 0.9338 | NS |
| Blood Hb concentration | NS | $F_{1,42}$ = 59.4601<br>P < 0.0001* | $F_{4,42}$ = 1.3807<br>P = 0.2745 | NS |
| Red blood cell $P_{50}$ | NS | $F_{1,42}$ = 18.8344<br>P = 0.0002* | $F_{4,42}$ = 3.9622<br>P = 0.0150* | NS |
| Hill coefficient | NS | $F_{1,42}$ = 0.0315<br>P = 0.8599 | $F_{4,42}$ = 0.2514<br>P = 0.9073 | NS |

*P < 0.05. [1]Interaction between acclimation and globin genotype. NS denotes no significant effect of the factor, which was removed from the final statistical model. [2]Mouse family was also included as a random factor in the mixed model, for which 0.05 < P < 0.1.

**Table S3.** Effects of inspired $PO_2$ and hemoglobin genotype on cardiorespiratory physiology of $F_2$ inter-population hybrid deer mice acclimated to normoxia.

| Trait | Animal mass | $PO_2$ | Hb genotype | Interaction[1] |
|---|---|---|---|---|
| $\dot{V}O_2max$ | $F_{1,41}$ = 31.2238<br>P < 0.0001* | $F_{1,41}$ = 61.0119<br>P < 0.0001* | $F_{4,41}$ = 3.0346<br>P = 0.0416* | NS |
| Arterial $O_2$ saturation | NS | $F_{1,41}$ = 122.1012<br>P < 0.0001* | $F_{4,41}$ = 3.7422<br>P = 0.0189* | $F_{4,41}$ = 3.0674<br>P = 0.0389* |
| Heart rate[2] | NS | $F_{1,41}$ = 32.4464<br>P < 0.0001* | $F_{4,41}$ = 2.8000<br>P = 0.0545 | NS |
| Total ventilation | NS | $F_{1,42}$ = 0.3248<br>P = 0.5738 | $F_{4,42}$ = 0.6799<br>P = 0.6136 | NS |
| Tidal volume | NS | $F_{1,42}$ = 0.1428<br>P = 0.7087 | $F_{4,42}$ = 1.4425<br>P = 0.2551 | NS |
| Breathing frequency | $F_{1,41}$ = 5.9183<br>P = 0.0242* | $F_{1,41}$ = 0.8443<br>P = 0.3669 | $F_{4,41}$ = 1.0643<br>P = 0.3999 | NS |
| Hematocrit | NS | NA | $F_{4,21}$ = 0.3604<br>P = 0.8339 | NA |
| Blood Hb concentration | NS | NA | $F_{4,21}$ = 0.2008<br>P = 0.9351 | NA |
| Red blood cell $P_{50}$ | NS | NA | $F_{4,21}$ = 5.1298<br>P = 0.0048* | NA |
| Hill coefficient | NS | NA | $F_{4,21}$ = 0.4016<br>P = 0.8053 | NA |

*P < 0.05. [1]Interaction between inspired $PO_2$ and globin genotype. NS denotes no significant effect of the factor, which was removed from the final statistical model. NA, not applicable.

[2]Mouse family was also included as a random factor in the mixed model, for which P < 0.05.

**Table S4.** Effects of acclimation to hypoxia and globin genotype on cardiorespiratory physiology in normoxia of $F_2$ inter-population hybrid deer mice.

| Trait | Animal mass | Acclimation | Hb genotype | Interaction[1] |
|---|---|---|---|---|
| Normoxic $\dot{V}O_2max$[2] | $F_{1,40} = 23.2664$<br>$P < 0.0001$* | $F_{1,40} = 4.2325$<br>$P = 0.0502$ | $F_{4,40} = 1.6212$<br>$P = 0.2115$ | NS |
| Arterial $O_2$ saturation | $F_{1,41} = 15.3290$<br>$P = 0.0003$* | $F_{1,41} = 0.4014$<br>$P = 0.5296$ | $F_{4,41} = 2.9739$<br>$P = 0.0291$* | NS |
| Heart rate[2] | $F_{1,40} = 4.8558$<br>$P = 0.0381$* | $F_{1,40} = 0.1977$<br>$P = 0.6605$ | $F_{4,40} = 2.6116$<br>$P = 0.0698$ | NS |
| Total ventilation | NS | $F_{1,42} = 22.9660$<br>$P < 0.0001$* | $F_{4,42} = 0.3536$<br>$P = 0.8386$ | NS |
| Tidal volume | $F_{1,41} = 5.1639$<br>$P = 0.0317$* | $F_{1,41} = 3.1282$<br>$P = 0.0892$ | $F_{4,41} = 1.5707$<br>$P = 0.2199$ | NS |
| Breathing frequency | NS | $F_{1,42} = 8.0995$<br>$P = 0.0087$* | $F_{4,42} = 0.6049$<br>$P = 0.6634$ | NS |

*$P < 0.05$. [1]Interaction between acclimation and globin genotype. NS denotes no significant effect of the factor, which was removed from the final statistical model. [2]Mouse family was also included as a random factor in the mixed model as $P < 0.05$.

**Table S5.** Parameters used to generate the initial solution in the model of the oxygen transport pathway representing the 'ancestral condition' with the most lowland $P_{50}$.

| Variable | Value |
|---|---|
| *Measured input parameters* | |
| $P_B$ (kPa) | 101 |
| $F_IO_2$ | 0.123 |
| $\dot{V}$ (ml min$^{-1}$ g$^{-1}$) | 4.96 |
| $V_T$ (µl g$^{-1}$) | 13.0 |
| [Hb] (g dl$^{-1}$) | 14.2 |
| $P_{50}$ (kPa) | 4.84 |
| *n* | 2.85 |
| $T_b$ (˚C) | 31.4 |
| *Estimated input parameters* | |
| $V_D$ (µl g$^{-1}$)* | 6.40 |
| $\dot{Q}$ (ml min$^{-1}$ g$^{-1}$) † | 1.06 |
| *Calculated input parameters* | |
| $D_LO_2$ (ml kPa$^{-1}$ min$^{-1}$) | 0.0661 |
| $D_TO_2$ (ml kPa$^{-1}$ min$^{-1}$) | 0.0322 |
| *Output parameters (ancestral values shown)* | |
| $\boldsymbol{P_AO_2}$ (kPa) | 7.18 |
| $\boldsymbol{P_aO_2}$ (kPa) † | 6.85 |
| $\boldsymbol{P_vO_2}$ (kPa)† | 2.29 |
| $\boldsymbol{\dot{V}O_2}$**max** (ml min$^{-1}$ g$^{-1}$) | 0.131 |

$P_B$, barometric pressure; $F_IO_2$, inspired oxygen fraction; $\dot{V}O_2$max, maximal oxygen consumption rate measured during acute cold exposure; $\dot{V}$, total ventilation; $V_T$, tidal volume; [Hb], blood hemoglobin concentration; $P_{50}$, $PO_2$ at 50% $O_2$ saturation; *n*, Hill coefficient; $P_aO_2$, arterial $O_2$ tension; $T_b$, body temperature; $V_D$, dead space volume; $\dot{Q}$, cardiac ouput; $P_vO_2$, mixed venous $O_2$ tension; $P_AO_2$, alveolar $O_2$ tension; $D_LO_2$, $O_2$ diffusing capacity of the lungs; $D_TO_2$, $O_2$ diffusing capacity of the tissues. *Indicates value was taken from Fallica *et al* 2011 (1). †Indicates value was taken from, or calculated using, data in Tate *et al* 2020 (2). Variables in bold were then calculated by the model in our sensitivity analysis in response to changes in $D_TO_2$ and/or $P_{50}$.

**References**

1. J. Fallica, S. Das, M. Horton, W. Mitzner, Application of carbon monoxide diffusing capacity in the mouse lung. *J. Appl. Physiol.* **110**, 1455-1459 (2011).

2. K. B. Tate *et al.*, Coordinated changes across the O2 transport pathway underlie adaptive increases in thermogenic capacity in high-altitude deer mice. *Proc. R. Soc. London., B, Biol. Sci.* **287**, 20192750 (2020).
